# Recapitulation and reversal of schizophrenia-related phenotypes in *Setd1a*-deficient mice

**DOI:** 10.1101/529701

**Authors:** Jun Mukai, Enrico Cannavò, Gregg W. Crabtree, Ziyi Sun, Anastasia Diamantopoulou, Pratibha Thakur, Chia-Yuan Chang, Yifei Cai, Stavros Lomvardas, Atsushi Takata, Bin Xu, Joseph A. Gogos

## Abstract

*SETD1A*, a lysine-methyltransferase, is a key schizophrenia susceptibility gene. Mice carrying a heterozygous loss-of-function mutation of the orthologous gene exhibit alterations in axonal branching and cortical synaptic dynamics, accompanied by working memory deficits. We show that Setd1a binds both promoters and enhancers with a striking overlap between Setd1a and Mef2 on enhancers. Setd1a targets are highly expressed in pyramidal neurons and display a complex pattern of transcriptional up- and down-regulations shaped by presumed opposing functions of Setd1a on promoters and Mef2-bound enhancers. Notably, evolutionary conserved Setd1a targets are associated with neuropsychiatric genetic risk burden. Reinstating *Setd1a* expression in adulthood rescues cognitive deficits. Finally, we identify LSD1 as a major counteracting demethylase for Setd1a, and show that its pharmacological antagonism results in a full rescue of the behavioral and morphological deficits in *Setd1a*-deficient mice. Our findings advance understanding of how *SETD1A* mutations predispose to SCZ and point to novel therapeutic interventions.

## Introduction

Schizophrenia (SCZ) is a disabling neuropsychiatric disorder defined by positive and negative symptoms as well as cognitive impairment. Over the past decade, human genetic studies have revealed an important role of individually rare (but collectively common) *de novo* (DNMs) and inherited mutations in genetically complex neuropsychiatric diseases (Fromer et al., 2014; Genovese et al., 2016; Gulsuner et al., 2013; Rodriguez-Murillo et al., 2012). We have previously linked loss-of-function (LoF) mutations in *SETD1A*, a histone methyltransferase, to risk for SCZ (Takata et al., 2016; Takata et al., 2014). This finding was subsequently confirmed by large scale exome sequencing studies (Singh et al., 2016; Takata et al., 2016; Takata et al., 2014). SETD1A mutations have also been reported in early-onset epilepsy (Yu et al., 2019) and disrupted speech development (Eising et al., 2019). SETD1A is part of the Set/COMPASS complex, which mediates mono-, di-, and tri-methylation of the lysine 4 on the histone H3 protein (H3K4) (Miller et al., 2001). In humans, Set/COMPASS is comprised of one of six catalytic subunits [SETD1A, SETD1B, KMT2A, KMT2D, KMT2C and KMT2B], four common subunits (WDR5, RBBP5, ASH2L, and DPY30), and several subtype-specific proteins (Shilatifard, 2012). In addition to *SETD1A*, mutations in subunits of Set/COMPASS have been reported in SCZ and other neurodevelopmental disorders (NDD) (Gulsuner et al., 2013; Neale et al., 2012; O’Roak et al., 2012) as well as dominant Mendelian syndromes (Fahrner and Bjornsson, 2014).

*SETD1A* is a confirmed SCZ risk gene and *SETD1A* mutations confer a large increase in disease risk (Singh et al., 2016), which provides an excellent genetic substrate for disease modeling. However, the impact of SETD1A deficiency on brain structure and function and the mechanisms by which a ubiquitous molecular process, such as histone methylation, leads into specific psychiatric disease symptoms remains unknown. Here, we performed a comprehensive analysis of mutant mice carrying a LoF allele in the endogenous *Setd1a* orthologue that models the various identified *SETD1A* SCZ LoF risk alleles to uncover the role of *SETD1A* in gene regulation, neuronal architecture, synaptic plasticity and behavioral paradigms dependent on learning. We also used this mouse model to examine whether malfunction of neural circuits and neurocognitive deficits emerging due to *Setd1a*-deficiency can be reversed by pharmacological interventions during adulthood, thus paving the way for the development of new therapeutics.

## Results

The Allen Brain Atlas (www.brain-map.org) predicts that *Setd1a* is expressed throughout the adult mouse brain, with higher expression in neocortex. Real-time Quantitative Reverse Transcription PCR (qRT-PCR) analysis on mouse prefrontal cortex (PFC) detected Setd1a mRNA at various developmental stages (Figure S1A). Immunostaining of the prelimbic area in medial PFC of 6-week old mice revealed that Setd1a positive cells are distributed in all cortical layers except layer 1 (L1) and merged with NeuN positive (NeuN^+^) cells (Figure S1B). Setd1a protein exhibits mixtures of punctate and diffuse fibrillary nuclear staining in all NeuN^+^ neuron and parvalbumin positive (PV^+^) interneurons in the medial PFC superficial layers, but only a few small puncta in CNPase (oligodendrocytes) and GFAP (astrocytes) positive cells in the cortical plate and corpus callosum (Figure S1C). Setd1a localizes to euchromatin and does not overlap with heterochromatin regions in either NeuN^+^ or PV^+^ neurons (Figure S1C).

We employed a mouse model carrying a *Setd1a* LoF allele (Figures S1D and S1E). Reduction of *Setd1a* RNA and protein levels to approximately half of WT levels was confirmed by qRT-PCR and immunoblots of adult frontal cortex (Figures S1F and S1G). Although *Setd1a^-/-^* is embryonic lethal (Bledau et al., 2014), *Setd1a^+/-^* mice were indistinguishable from WT littermates in terms of body size, weight and posture (not shown).

### *Setd1a^+/-^* mice exhibit deficits in working memory (WM) and cortical synaptic plasticity

Cognitive impairment is a core feature of SCZ (Barch and Ceaser, 2012). Histone methylation mechanisms play a role in the coordination of complex cognitive processes and in modulating the genetic risk and the phenotypic variation of cognitive disorders (Rudenko and Tsai, 2014). To evaluate the relative effects of *Setd1a* deficiency on different cognitive domains and underlying brain circuits, we first asked whether *Setd1a* deficiency in mice affects performance in a variety of cognitive tasks. First, we established that *Setd1a^+/-^* mice show normal activity in the open-field assay (Figures S1H and S1I). We assessed spatial WM performance using a T-maze delayed non-match to place task, in which a mouse is required to remember a reward location over a delay and following the delay it is reinforced for making a response towards an alternative location (see Methods). *Setd1a^+/-^* mice learned the task and performed almost as well as WT littermates during training (Figure 1A and 1B). In the subsequent WM test, however, *Setd1a^+/-^* mice manifested a marked impairment (correct responses: 63.63 ± 3.03 %, *P* = 0.0038 at 10 sec; 64.58 ± 3.10 %, *P* = 0.0172 at 30 sec), compared to WT littermate controls (76.39 ± 2.68 % at 10 sec, 76.39 ± 3.38 % at 30 sec) (Figure 1C). In contrast, memory performance of *Setd1a^+/-^* mice was intact in a series of other cognitive tasks, including direct interaction test for social memory (Figure S1J), novel object recognition test (Figure S1K) and fear conditioning task for associative memory (Figure S1L). Thus, *Setd1a* deficiency leads to robust but circumscribed cognitive impairments in spatial WM, a core cognitive phenotype of psychotic disorders (Arguello and Gogos, 2012).

**Figure 1.**
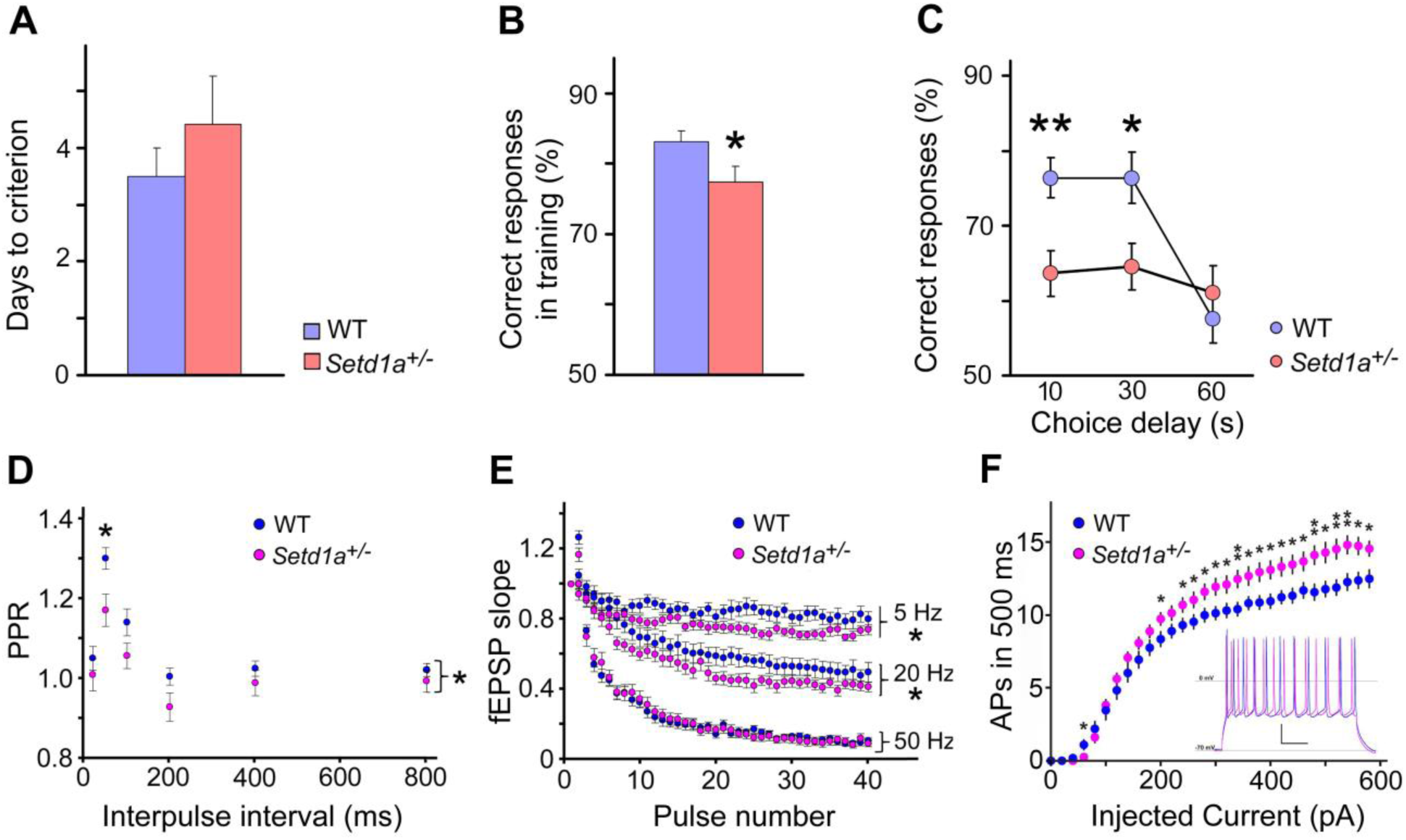
*Setd1a* deficiency impairs WM and alters short-term plasticity and neuronal excitability in PFC. (A) Days taken to reach criterion performance (three consecutive days of > 70% choice accuracy) on the delayed non-match to sample T-maze task. *P* = 0.36. N = 12 each genotype. (B) Correct responses during training on the delayed non-match to sample T-maze task. *P* = 0.041. N = 12 each genotype. (C) Working memory performance in a delayed non-match to place task. Mean percentage of correct responses for WT and *Setd1a^+/-^* mice (N = 12 each genotype). Note that vertical axis starts at 50% correct responses, which represents baseline response accuracy expected by chance. *P* = 0.0038 (10 sec), 0.0172 (30 sec), 0.485 (60 sec). N, number of independent animals. Data are shown as means ± s.e.m. \**P* < 0.05, \*\**P* < 0.01. Student’s two-tailed *t-*test (A, B, C). (D and E) *Setd1a* deficiency impairs short-term plasticity in PFC. (D) PPR is significantly decreased in *Setd1a*^+/-^ mice (N = 7, n = 23) at all tested ISIs (20, 50, 100, 200, 400 and 800 ms), compared to WT mice (N = 7, n = 23) (2-way repeated-measures ANOVA, *P* < 0.05; Bonferroni posttests revealed a significant difference at ISI of 50 ms, *P* < 0.05). (E) Frequency-dependent differences of STD of the fEPSPs in WT and *Setd1a*^+/-^ mice at 5 Hz, 20 Hz and 50 Hz. At 5 and 20 Hz, the STD is significantly greater in *Setd1a*^+/-^ mice (N = 7, n = 22 at 5 Hz and N = 7, n = 19 at 20 Hz) compared to their WT controls (N = 7, n = 23 at 5 Hz and N = 7, n = 20 at 20 Hz; 2-way repeated-measures ANOVA, *P* < 0.05 for time × genotype interaction). At 50 Hz, the STD is similar between genotypes across experiments (WT mice: N = 7, n = 18 and *Setd1a*^+/-^ mice: N = 7, n = 16; 2-way repeated-measures ANOVA, *P* > 0.05). N, number of independent animals. n, number of brain slices. Data are shown as means ± s.e.m. \**P* < 0.05, \*\**P* < 0.01. (F) Summary data (mean ± s.e.m) of number APs evoked in response to 500 ms currents steps. WT recordings, blue; *Setd1a*^+/-^ recordings, magenta. (WT, n = 22, *Setd1a*^+/-^, n = 20; 2-way RM ANOVA, *P* = 0.021). Values represent mean ± s.e.m. (Starred data represent results of pair-wise t-test comparisons at given current steps: * *P <* 0.05, ** *P <* 0.01). *Inset; Setd1a*^+/-^ neurons display enhanced excitability (magenta trace) compared to WT neurons (blue trace). Representative traces from current-clamp recordings from layer 2/3 pyramidal neurons showing AP responses to near-maximal current step (500 pA, 500 ms). Scale bar: 20 mV, 100 ms. Reference levels, V_rest_ = -70 mV, and 0 mV.

WM is sensitive to PFC dysfunction and can be affected by the synaptic properties of PFC neurons (Barak and Tsodyks, 2014). The glutamatergic inputs from L2 neurons to the apical and basal dendrites of L5 pyramidal neurons form monosynaptic synapses in medial PFC, which are critical for controlling the computations and output of L5 pyramidal neurons (Quiquempoix et al., 2018) and play a key role in WM and goal directed behaviors (Bastos et al., 2018; Goldman-Rakic, 1995). We performed field recordings in brain slices on both genotypes to examine the synaptic properties in L5 neurons in response to the stimulation of L2. The input-output relation across experiments was comparable between genotypes, which indicates normal basic synaptic transmission. Similarly, the fiber volley amplitude was unaltered in *Setd1a^+/-^* mice indicating that when the same number of afferent fibers is activated, basal transmission is unaffected (Figures S2A, S2B, and S2C). We next determined whether short-term synaptic plasticity was impaired in *Setd1a*^+/-^ mice. To this end, we employed a paired-pulse ratio (PPR) protocol at various inter-stimulus intervals (ISIs) as previously described for two other genetic models (Crabtree et al., 2017; Fenelon et al., 2013). At ISIs of less than 200 ms, both genotypes exhibited a significant facilitation, but *Setd1a*^+/-^ mice showed overall significantly decreased PPR values (Figures 1G and S2J, N = 7, n = 23; *Setd1A*^+/-^, N = 7, n = 23; two-way repeated-measures ANOVA, *P* = 0.04. Bonferroni post-tests further revealed a difference at ISI of 50 ms, *P* < 0.05) indicating an increased initial probability of presynaptic neurotransmitter release.

During PFC-dependent WM tasks, PFC neurons in local ensembles receive trains of inputs from neighboring cells in the 20 to 60 Hz frequency range (Miller et al., 1996). We tested the effect of *Setd1a* deficiency on synaptic plasticity within this physiological frequency range by applying 40 stimuli at 5 Hz, 20 Hz and 50 Hz. In response to these stimulus trains, a rapid depression of synaptic strength was observed in both genotypes, attributable to depletion of the readily releasable vesicle pool (Zucker and Regehr, 2002). Compared to WT mice, *Setd1a*^+/-^ mice show significantly greater depression of fEPSP responses at 5 Hz (Figures 1H and S2K, WT, N = 7, n = 23; *Setd1a*^+/-^, N = 7, n = 22; two-way repeated-measures ANOVA, *P* = 0.03) and at 20 Hz (Figures 1H and S2K, WT, N = 7, n = 20; *Setd1a* ^+/-^, N = 7, n = 19; two-way repeated-measures ANOVA, *P* = 0.04). No genotypic difference was observed at 50 Hz (Figures 1H and S2K, WT, N = 7, n = 18; *Setd1A*^+/-^, N = 7, n = 16; two-way repeated-measures ANOVA, *P* = 0.95). To determine the impact of *Setd1a* deficiency on the potentiation of synaptic efficacy of L5 cortical synapses, short-term (STP) and long-term potentiation (LTP) were induced as described previously (Crabtree et al., 2017; Fenelon et al., 2013). No differences were found in either one of these measures (Figures S2D and S2L, *P* = 0.57).

In addition to synaptic function, we tested whether PFC neurons displayed alterations in intrinsic excitability. Neuronal excitability is known to critically regulate network information flow (Titley et al., 2017) and has previously been shown to be altered in animal models of SCZ (Benamer et al., 2018; Crabtree et al., 2017; Gruter et al., 2015; Hamm et al., 2017). To test whether *Setd1a*-deficient neurons displayed altered intrinsic excitability, we performed whole-cell current-clamp recordings in PFC L2/3 pyramidal neurons – a major source of excitatory input to L5 – and found significantly increased action potential (AP) responses to current steps (Figure 1I). In contrast, we found the minimal current required for AP generation (rheobase) was unaltered (Figure S2E). Notably, neuronal input resistance, membrane time constant, and resting membrane potential were all found to be unaltered and thus did not underlie the observed enhanced neuronal excitability (Figures S2F, S2G and S2H). Both initial and terminal inter-spike intervals were significantly reduced (Figure S2M) while AP afterhyperpolarization was not altered (Fig S2N). Subsequent whole-cell voltage-clamp studies employing voltage steps revealed that *Setd1a^+/-^* mice displayed reduced outward currents near AP threshold voltages (-40mV to -30mV, Figures S2I and S2O). These reductions in hyperpolarizing currents may in part explain the enhanced AP generation we observed in *Setd1a^+/-^* mice.

### Laminar organization and cytoarchitecture in the cortex of *Setd1a^+/-^* mice

Adult 8-week-old *Setd1a^+/-^* mice manifest normal gross brain morphology and only a modestly diminished overall density of neurons (as identified by their NeuN immunoreactivity) in L2/3 and L5. Quantitative analysis of different interneuron subpopulations as identified by their immunohistochemical markers revealed a reduced number of PV^+^ inhibitory neurons in L5 and Reelin^+^ inhibitory neurons in L2/3 and L5, in the prelimbic area of medial PFC (Figures S3A, S3B, S3C, S3E, and S3H), while the numbers of VIP and SST positive interneurons were unchanged (Figures S3D, S3G, S3F, and S3). Further analysis (Figures S3J, S3K, S3L, and S3M) showed that alterations in the laminar organization are, at least in part, due to alterations in the proliferation of neural progenitor cells (NPCs) during corticogenesis, as also previously suggested (Bledau et al., 2014). Whether alterations in the numbers of interneuron populations also reflect homeostatic responses to altered circuit activity remains to be determined.

We also investigated whether *Setd1a* deficiency affects development of cortical axons, dendrites and spines. Plasmids encoding EGFP were electroporated *in utero* into the dorsolateral ventricular zone of the mouse forebrain at E15.5, and embryos were allowed to be born and develop to P8.5 for analysis. All cells labeled with EGFP completed their migration into L2/3 by P8.5. As shown previously (Mukai et al., 2015), callosal axons crossed the midline, coursed beneath the gray matter of the contralateral somatosensory cortex, penetrated the gray matter, and ascended to L1–3 and 5 (Figures 2A and 2B). The total number of callosal axon-terminal branches within the contralateral cortical L1–4 was decreased by disruption of *Setd1a* (7.86 ± 0.46, N = 10, n = 30, *P* < 0.0001), compared to WT brains (12.20 ± 0.44, N = 10, n = 30) (Figures 2C and 2D). Notably secondary branch number was robustly decreased in *Setd1a^+/-^* brains (1.95 ± 0.44, N = 10, n = 30 versus WT brains 3.63 ± 0.48, N = 10, n = 30; *P* < 0.007) (Figures 2C and 2D). Similar deficits in axonal branching were observed in cortical primary cultures from *Setd1a^+/-^* and WT mice (Figures 2E, 2F, 2G, 2H, and 2I).

**Figure 2.**
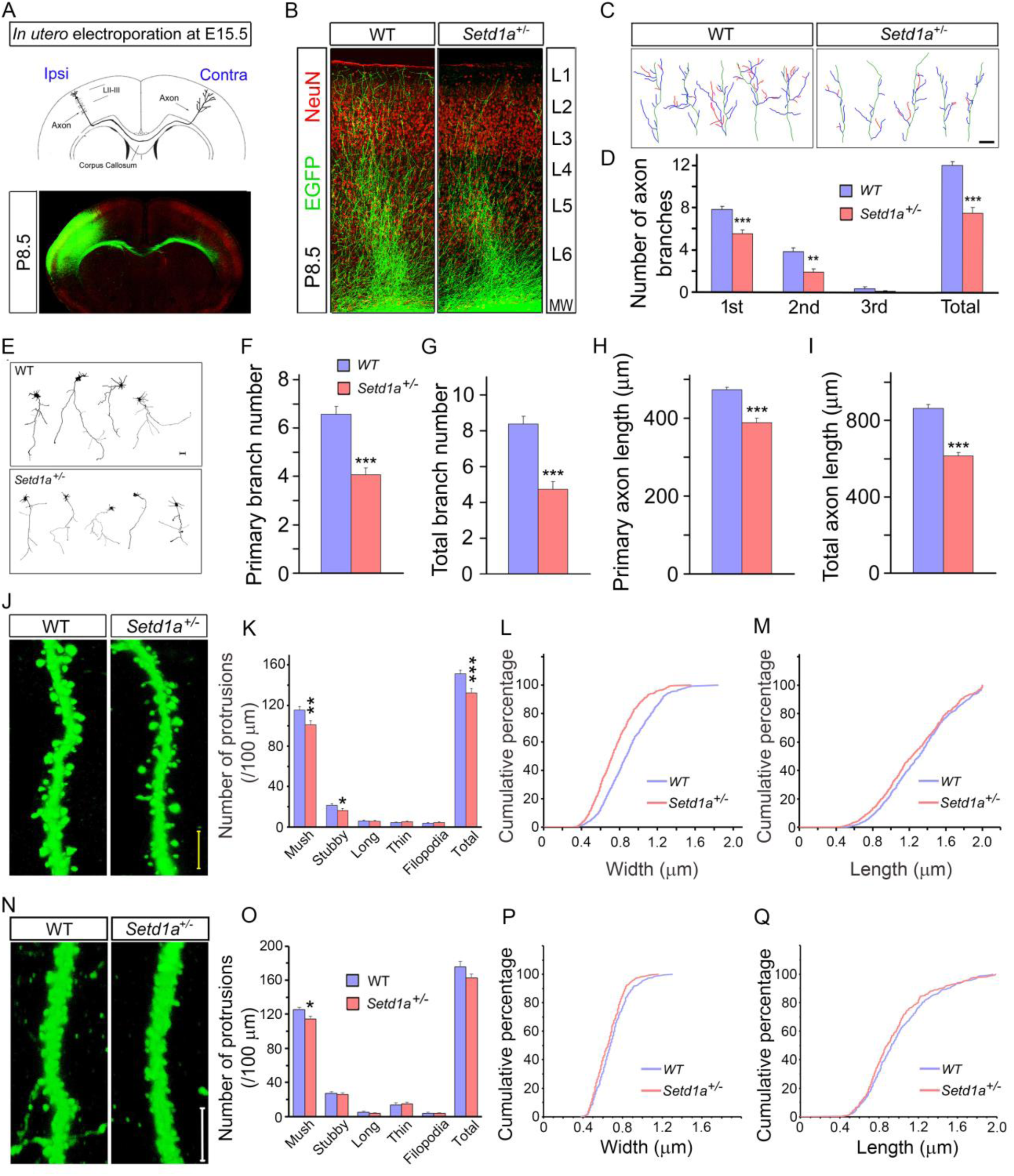
*Setd1a* deficiency disrupts axonal growth and spine morphology. (A-D) *Setd1a* deficiency disrupts axonal terminal branching *in vivo*. (A) Illustration of the callosal axon pathway of neurons residing at L2/3, electroporated with EGFP at E15.5 and assayed at P8.5 (*upper panel*). Lower panel shows EGFP-labeled neurons (*green*) and NeuN (*red*). Representative images (*lower panel*) are composites of more than one image acquired from one brain section under identical scanning parameters. (B) Representative images of contralateral axon terminal branching of EGFP-labeled neurons from coronal sections of P8.5 brains of mice *in utero* electroporated at E15.5. Images are composites of more than one image acquired from one brain section under identical scanning parameters. (C) Tracings of representative neurons in *in utero* electroporated brains expressing EGFP at P8.5, from similar sections as in (B). Depicted are 2D projections of the axon terminal in the contralateral cortical L1–4. Primary (*blue*), secondary (*red*), and tertiary (*pink*) branches from axons (*green*) are highlighted. Scale bar, 20 mm. (D) Quantitative assessment of contralateral axon branching in L1–4 reveals a reduction of branch number in *Setd1a^+/-^* mice (N = 10, n = 30 each genotype). (E-I) *Setd1a* deficiency disrupts axonal growth *in vitro*. (E) Representative images of axonal deficits in *Setd1a^+/-^* cortical neurons expressing EGFP at DIV5. *Setd1a^+/-^* cortical neurons were transfected with plasmids expressing EGFP at DIV4 and fixed at DIV5. Scale bar, 50 mm. (F) Quantification of primary branchpoint number. *Setd1a* deficiency caused a decrease in the number of primary branch points in *Setd1a^+/-^* neurons (4.1 ± 0.28, N = 4, n = 315, *P* < 0.001) compared to WT neurons (6.6 ± 0.31, N = 4, n = 304). (G) Quantification of total branchpoint number. Total branch points were reduced in *Setd1a^+/-^* (4.7 ± 0.68, N = 4, n = 315, *P* < 0.0009) compared to WT neurons (total branchpoints: 8.4 ± 0.76, N = 4, n = 304). (H) Quantification of primary axon length. Primary axon length was reduced at DIV5 in *Setd1a^+/-^* (388.6 ± 11.8 mm, N = 4, n = 315, *P* < 0.0008) compared to WT neurons (472.9 ± 6.9 mm, N = 4, n = 304). (I) Quantification of total axon length. Total axon length were reduced in *Setd1a^+/-^* (total axon length: 614.2 ± 22.6 mm, N = 4, n = 315, *P* < 0.0003) compared to WT neurons (total axon length: 869.6 ± 25.9 mm, N = 4, n = 304). (J) Alterations in spines of basal dendrites of L5 pyramidal neurons at the prelimbic area of medial PFC in *Setd1a^+/-^* mice. Representative, high-magnification images of eGFP–expressing neurons from WT Thy1-GFP/M (*left panel*) and *Setd1a^+/-^*; *Thy1-GFP/M* (*right panel*) mice. (K) Distribution of spine morphotypes, as well as total protrusions, in basal dendrites of L5 pyramidal neurons in prelimbic area from *Thy1-GFP^+/-^;Setd1a^+/-^* mice (N = 10, n = 20 each genotype). Dendritic protrusions were classified as different spine categories (mushroom, long, thin, or stubby) or as filopodia, and counted. Scale bar, 5 μm. (L and M) Reduction in the width (L; *P* < 0.0001, Kolmogorov-Smirnov test), and length (M; *P* < 0.001, Kolmogorov-Smirnov test) of mushroom spines of *Setd1a^+/-^*; Thy1-GFP^+/–^ mPFC pyramidal neurons relative to WT Thy1-GFP^+/–^ mPFC neurons. (L, M: WT, N = 10 mice, n = 644, *Setd1a^+/-^*, N = 10, n = 605). (N) Alterations in spines of basal dendrites of L2/3 pyramidal neurons in the prelimbic area of medial PFC in *Setd1a^+/-^* mice. Representative, high-magnification images of EGFP–expressing neurons from WT (*left panel*) and *Setd1a^+/-^* (*right panel*) mice (N), injected with AAV2-hSyn-EGFP virus. (O) Density of mushroom spines estimated over 100 μm of dendritic length in basal dendrites of L2/3 pyramidal neurons in the prelimbic area of medial PFC is modestly reduced in *Setd1a^+/-^* mice (N = 7, n = 21 each genotype). Scale bar, 5 μm. (P and Q) Reduction in the width (P) (*P* < 0.034, Kolmogorov-Smirnov test), but not length (Q) (*P* = 0.259, Kolmogorov-Smirnov test) of mushroom spines of *Setd1a^+/-^* L2/3 mPFC pyramidal neurons relative to WT neurons (P, Q: WT, N = 9 mice, n = 379, *Setd1a^+/-^*, N = 8, n = 373). \**P* < 0.05; \*\**P* < 0.01; \*\*\**P* < 0.001. Student’s two-tailed *t*-test (D, F-I, K, O). N: number of independent animals, n: number of neurons or brain slices. Histograms show means and error bars represent s.e.m.

We also looked for deficits in dendrite and spine development in the basal dendrites of L5 neurons in the prelimbic area of medial PFC by intercrossing *Setd1a^+/-^* mice with a reporter Thy1-GFP/M line, where L5 pyramidal neurons are labeled sparsely (Figure S3N). *Setd1a^+/-^* mice did not show a simplification in the dendritic arbors by Sholl analysis (Figures S3O, S3P, and S3Q). Morphotypic analysis of spines in 8-week-old mice showed that mushroom spine density was modestly reduced in the basal dendrites of L5 *Setd1a^+/-^* (100.0 ± 3.4 / 100 μm, N = 10, n = 20, *P* < 0.002) compared to WT neurons (118.3 ± 3.6/100 μm, N = 10, n = 20; Figures 2J and 2K). Mushroom spines showed a small decrease in the head width (N = 10, n = 605, Kolmogorov-Smirnov test: *P* < 0.0001) and length (N = 10, n = 605, Kolmogorov-Smirnov test: *P* < 0.02) in *Setd1a^+/-^* compared to WT neurons (N = 10, n = 644) (Figures 2L and 2M). We examined whether the observed alteration in spine development extend to other cortical layers. Analysis at the basal dendrites of L2/3 neurons of *Setd1a^+/-^* and WT littermates, labeled with EGFP via AAV2-Syn-EGFP virus injection into the prelimbic area of medial PFC, revealed a modest reduction is mushroom spine density (114.6 ± 3.4/100 μm, N = 7, n = 21 versus 125.4 ± 2.9 / 100 μm, N = 7, n = 21, *P* < 0.035; Figures 2N and 2O). Reduction in head width (N = 8, n = 373, Kolmogorov-Smirnov test: *P* < 0.034), but not length (*P* = 0.259) was also observed in *Setd1a^+/-^* L2/3neurons compared to WT neurons (N = 9, n = 379) (Figures 2P and 2Q).

### Identification of direct targets of Setd1a in the PFC of adult mice

Given that Setd1a is a lysine demethylase that regulates transcription, we sought to identify direct genomic targets of Setd1a in the PFC by chromatin immunoprecipitation followed by sequencing (ChIP–Seq, see Methods). We performed ChIP-Seq analysis in chromatin prepared from micro-dissected PFCs of 6-week old mice for histone modifications that mark active promoters (H3K4me2, H3K4me3 and H3K27ac) and active or poised enhancers (H3K4me1 and H3K27ac) using the same chromatin preparations. Using stringent criteria [Irreproducibility Discovery Rate (IDR) < 0.02], we identified 25,658 Setd1a peaks genome-wide. Analysis of the genome-wide binding profile suggests that, despite the fact that Setd1a mediates the trimethylation of H3K4 (Shilatifard, 2012), only a minority (34%, 8,769/25,658) of Setd1a peaks are located at promoter elements as observed by the overlap of Setd1a with H3K4me3 peaks (Figures 3A, 3B, S4A, and Data Table 1). The majority of Setd1a peaks (66%, 16,889/25,658) overlap with enhancer marks and not H3K4me3 (Figures 3B, S4A, Data Table 1). Further analysis of the genomic distribution of the non-promoter Setd1a peaks, revealed that 45% (7,601/16,889) reside within intergenic regions while 51% (8,567/16,889) reside in intronic sites with enhancer-like chromatin properties (Figures 3B, S4A). A minority resides within exons, 3’UTRs or within 1 kb from the transcription end-point (4%, 721/16,889).

**Figure 3.**
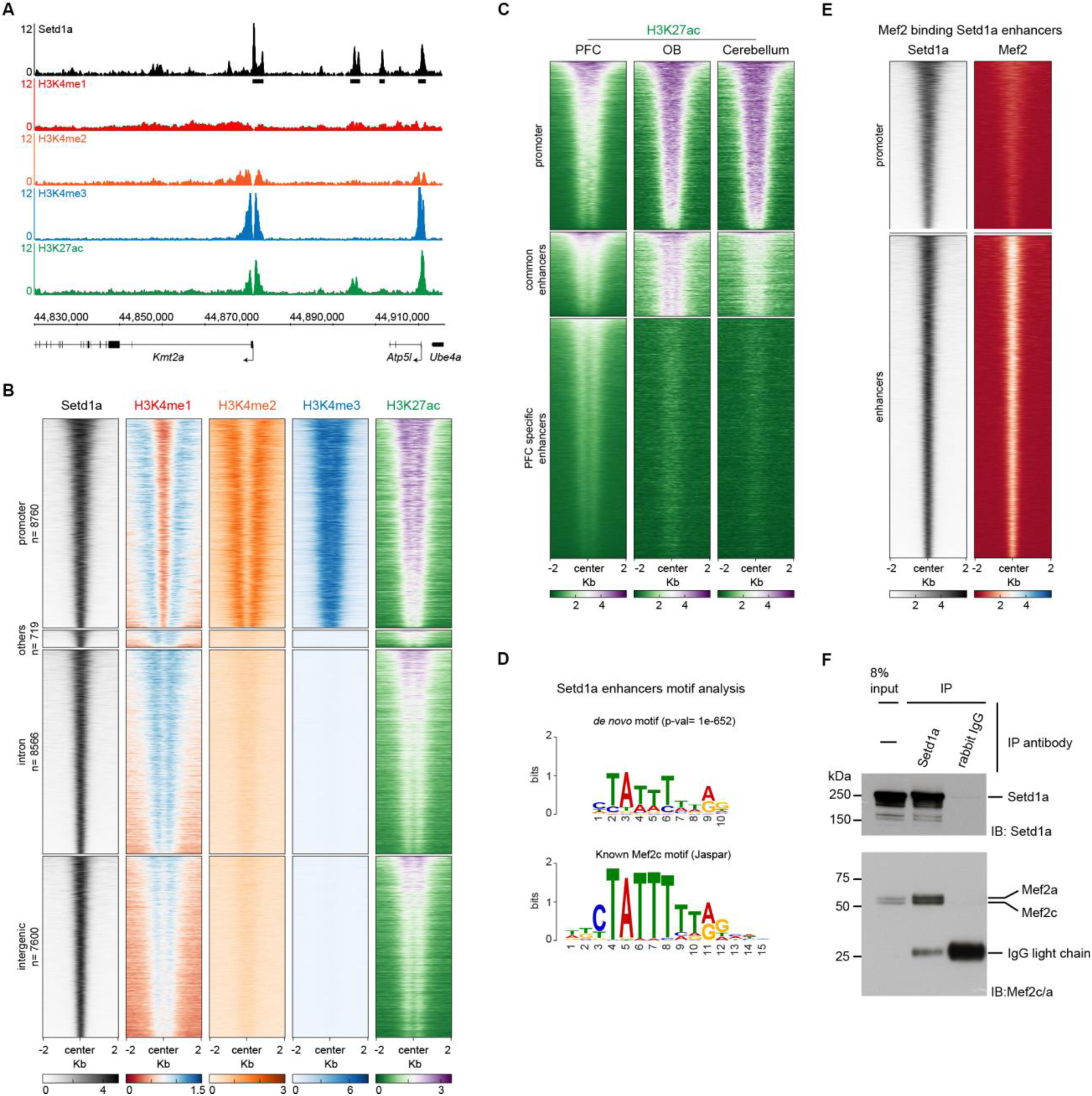
Setd1a binds promoters and enhancers in a complex with Mef2. (A) Representative ChIP-Seq locus showing Setd1a binding the promoter of *Kmt2a* gene. Normalized reads per genomic coverage from PFC ChIP-Seq for Setd1a (black) and chromatin marks (H3K4me1, red; H3K4me2, orange; H3K4me3, blue; H3K27ac, green) are shown. Black boxes below the Setd1a track represent significant peaks that passed all quality checks (see methods). (B) Heatmap of listed ChIP-Seq coverage from PFC, 2 kilobases upstream and downstream of the center of Setd1a peaks (n= 25,658). Setd1a peaks were clustered in promoters (*top panel*, defined by the overlap with H3K4me3 peaks) or enhancers (defined by absence of H3K4me3 signal and presence of H3K27ac and H3K4me1 signal) (n=16,889). Enhancers were sub-clustered by the annotation of Setd1a peaks. *Top-middle*, peaks in annotated 5′UTRs, coding sequence, transcription end site and 3′UTRs (defined as “others”) (n=721); *middle-bottom*, peaks in annotated introns (n=8567); *bottom panel*, peaks in intergenic regions (n=7601). All heatmaps are sorted in the same order, based on Setd1a and H3K27ac signal. (C) Heatmap of H3K27ac 2 Kb around Setd1a peaks in PFC, OB and Cerebellum. By clustering, we distinguished 3 different clusters depending on H3K27ac coverage across tissues. While promoter and a subset of enhancers marked as “common enhancers” (n= 4390) showed H3K27ac in all three analyzed tissues, we identified a third cluster of enhancers, marked as “PFC specific enhancers” (n=12499), that showed H3K27ac signal only in the PFC. (D) *De novo* transcription factor motif analysis in Setd1a bound promoters (*top panel*) and enhancers (*bottom panel*). Setd1a bound promoters motif analysis showed enrichment (*p-val*= 1e-142) for a motif very similar to the known Sp1 motif (bottom, Jaspar Core Vertebrata database). Instead, Setd1a bound enhancers showed enrichment (*p-val*= 1e-852) for the known Mef2c motif (Jaspar Core Vertebrata database). (E) Heatmap of Setd1a and Mef2 coverage 2 kb around the center of Setd1a peaks in promoters (*top panel*) and enhancers (*bottom panel*). (F) Co-IP of Setd1a and Mef2a/c from the cortical plate of 6 week old mice. Western blot of 8% input (*left lane*), immunoprecipitation (IP) with Setd1a antibody (*middle lane*) or IgG (*right lane*) is shown. Detection with Mef2a/c, Setd1a and IgG antibody is shown in methods.

The *Setd1a^+/-^* phenotypic profile suggest a prominent impact of *Setd1a* deficiency on cortical structure and function. To assess if Setd1a-bound regulatory elements have pan-neuronal or cortical-specific functions we compared our H3K27ac ChIP-Seq from the PFC with ENCODE H3K27ac ChIP-Seq from the olfactory bulb (OB) and the cerebellum (Shen et al., 2012). Setd1a bound PFC promoters, as a whole, have H3K27ac in the OB and less so in the cerebellum (Figure 3C). In contrast the majority of the enhancer peaks (74%, 12,499/16,889) are enriched for H3K27ac specifically in the PFC (Figures 3C). These observations suggest that cortex-specific transcription factors recruit Setd1a at enhancers whereas more generally expressed transcription factors recruit Setd1a to promoters.

### Setd1a targets are extensively enriched for Mef2 binding sites

*De novo* motif analysis of the Setd1a peaks using HOMER revealed Mef2 consensus motifs as the most enriched DNA motif within the 16,889 enhancer peaks (Figure 3D, Data Table 1) and Sp1 consensus motifs as the most enriched motifs on the promoters (Figure 3D, Data Table 1). ChIP-Seq experiments with an antibody that recognizes Mef2a, Mef2c and Mef2d isoforms, which are expressed in the PFC, showed a striking overlap between Setd1a and Mef2 targets specifically at enhancers, with 11,688 of the 16,889 Setd1a peaks being occupied by Mef2 (Figure 3E S4B). Co-immunoprecipitation experiments with nuclear extracts prepared from the cortical plate, revealed an interaction between Setd1a and Mef2a/c (Figures 3F). To determine if this regulatory relationship is conserved across evolution, we examined the PhastCons conservation score (see methods) in the Euarchontoglire clade of Setd1a bound enhancers. This analysis revealed both conserved (∼46%, 7,738/16,889) and non-conserved (∼54%, 9,151/16,889) Setd1a targets. Among the former, we detected conservation of Mef2 binding sites, providing additional support for a functional role of Mef2 in Setd1a recruitment (Figure S4C and S4D). To further investigate if the interaction between Setd1a and Mef2 is conserved in humans in disease-relevant brain areas, we mapped Setd1a peaks from the mouse genome (mm10) to the corresponding positions in the human genome (hg38) focusing in particular to PFC-specific bound regions. We mapped 3,813 mouse genomic regions (out of 12,499) to the human genome (Data Table 1). To enrich for functional elements, we explored the chromatin accessibility of these peaks in the human brain. To this end, we analyzed a previously published ATAC-Seq dataset from 14 regions of human postmortem brains (Fullard et al., 2018) focusing our attention specifically on neuronal and non-neuronal cells of 5 brain areas: dorsal and ventral PFC, hippocampus, medial dorsal thalamus and putamen. Of the 3,813 conserved human peaks, 1,765 encompass significant ATAC-Seq peaks in at least one of the considered brain areas. Interestingly, we observed that these sites are accessible in neuronal cell types and not glia. Moreover, we found that a minority of peaks is accessible across all five regions of the brain (14%, 243/1,765) while the majority is mostly accessible in cortical regions and hippocampus (86%, 1,522/1,765) (Figure S4E). To further investigate the functionality of these sites in the human neurons, we performed motif analysis on the accessible conserved Setd1a regions with HOMER and identified MEF2 motifs as one of the most enriched motifs in the orthologous human sequences (*P* = 1e-41; Data Table 1). Altogether, these data suggest that the role of Mef2 and Setd1a in transcriptional regulation is conserved between mouse and human and that the intergenic Setd1a peaks we detected likely originate from neuronal cortical populations (see also below).

### Setd1a targets are enriched for orthologues of disease-risk genes

To elucidate whether Setd1a-dependent genome regulation in mice is linked to neuropsychiatric etiology in humans, we determined more directly if conserved Setd1a genomic binding sites are enriched for genes and genomic intervals implicated in SCZ by human GWAS studies. We first performed an interval-based analysis to assess enrichment of Setd1a peaks of the 108 loci identified by SCZ-GWAS (Schizophrenia Working Group of the Psychiatric Genomics, 2014). We focused only on PFC-specific enhancer peaks which are conserved and accessible in the human brain (see above) and applied a permutation test (N=20,000 permutations) implemented in the RegioneR program to determine if there is any significant overlap between these Setd1a peak intervals and the 108 SCZ-GWAS associated intervals (Schizophrenia Working Group of the Psychiatric Genomics, 2014). We found a significant overlap (*P* = 0.0055 Z =2.9) (Figures 4A and Data Table 2). This was true specifically for intronic Setd1a peaks (see Figure S5A-S5D). To confirm the specificity of this finding, we conducted an identical analysis on data from other GWAS of CNS and non-CNS diseases, with similar sizes of genome-wide significant intervals (Welter et al., 2014), but we did not observe any significant overlap (Tables S1 and S2).

**Figure 4.**
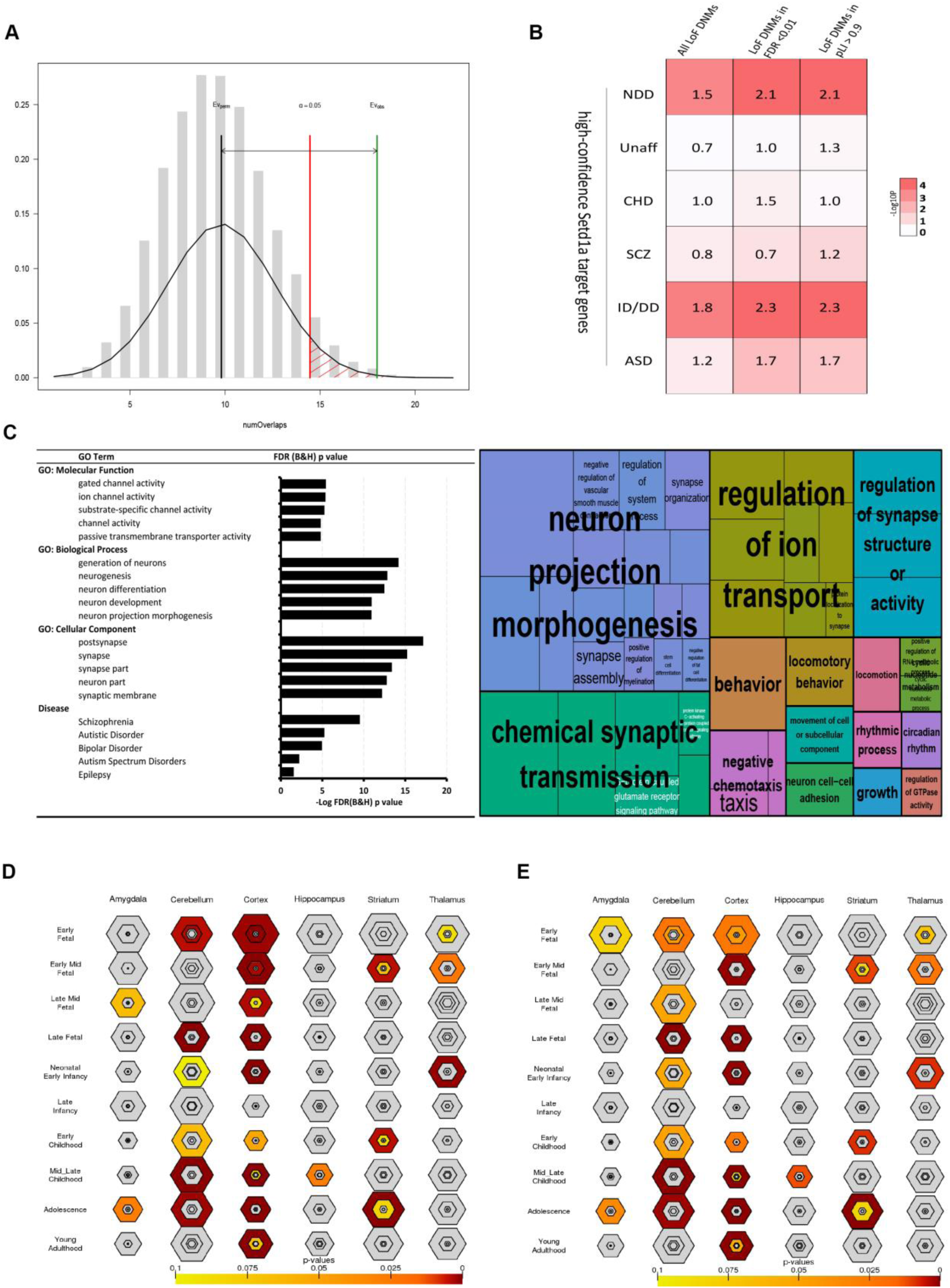
Setd1a targets are enriched for synaptic genes and overlap with genome-wide significant signals of SCZ and NDD. (A) Permutation tests (n=20,000, randomization method= circularRandomizeRegions) indicate there is significant overlaps between conserved and chromatin accessible Setd1a ChIP-Seq peaks (*P* = 0.0055, Z-score = 2.9) and the SCZ-GWAS loci identified by reference (Schizophrenia Working Group of the Psychiatric Genomics, 2014). X-axis is the number of overlaps, Y-axis is the density of expected number of overlaps determined by permutation. EV_perm_, Expected average number of overlaps by permutation, EV_obs_, actual observed number of overlaps. Permutation *P* value threshold = 0.05. (B) DNENRICH analysis of the high-confidence Setd1a target genes. Fold changes (observed/expected number of LoF DNMs in the gene-set of interest) are shown in the cells color-coded by p values. Unaff: Unaffected subjects, CHD: congenital heart diseases, SCZ: schizophrenia, ID/DD: intellectual disability/developmental disorders, ASD: autism spectrum disorders, NDD: neurodevelopmental disorders (SCZ+ID/DD+ASD) [FC; *P* (pLI/LoF-FDR): ID/DD: 2.3/2.3; 1.0 × 10^-6^ /1.0 × 10^-6^, ASD: 1.7/1.7; 0.002/0.005 and SCZ: 2.3/3.1; 0.065/0.022, total N of LoF DNMs in ID/DD, ASD and SCZ = 986, 793 and 95, respectively]. (C) GO-term analysis of target genes with all conserved accessible Setd1a ChIP-Seq peaks annotated by PAVIS and visualization of functional clusters using REVIGO GO treemap (Supek et al., 2011) for the enriched GO terms annotated by Toppgene. (D) Spatiotemporal expression pattern of target genes within conserved and accessible Setd1a peaks in human brain. *P* values are PSI *P* values as defined in the CSEA tool. (E) Spatiotemporal expression pattern of target genes within conserved and accessible Setd1a peaks that do not overlap with the mutation intolerant genes hit by LoF DNMs in NDD (as defined in B).

We followed this finding by a complementary gene-based analysis, which directly assess whether the observed enrichment of Setd1a regulatory elements within SCZ-GWAS loci extends to Setd1a target gene content, as well. We used the same set of conserved and accessible human enhancer peaks and restricted our analysis to targets genes with Setd1a peaks that represent well-annotated, high-confidence targets (all peaks with an annotation by PAVIS, N = 708 genes) as opposed to multiple ambiguous target genes abutting each intergenic peak (Data Table 2). We assessed enrichment of these high-confidence Setd1a targets within the SCZ-GWAS loci, using INRICH, and found a significantly overlap (FDR corrected *P* = 0.008, Table S3). The list of genes spans diverse neuronal functions ranging from neurogenesis to synaptic function and neuronal excitability and includes several putative risk-genes that have been identified in previous genetic studies of SCZ as well as other psychiatric and neurodevelopmental disorders including DPYD, TCF4, IMMP2L, DGKI, CSMD1, CACNB2, SATB2, CNTN4, GPM6A and GRM3 (Table S4).

Identical analysis of data from other GWAS of CNS and non-CNS diseases did not show significant enrichment except for Autism (Tables S3 and S5). Notably, in keeping with the hypothesis that Mef2 and Setd1a have a coordinated role in SCZ-related gene regulation we also observed an excess within SCZ-GWAS intervals of target genes with peaks co-bound by Setd1a and Mef2 but less significant with Mef2-only peaks (*P* = 0.003 and 0.03 respectively) (Table S6). We also investigated whether the high-confidence Setd1a target genes (N = 708) are significantly overrepresented among genes implicated in SCZ and other NDDs by studies of rare LoF DNMs. Using DNENRICH, we observed that there is significant enrichment of the high-confidence Setd1a target genes (Figure 4B, *P* = 1.6 × 10^-5^, FC = 1.5) among the genes hit by LoF DNMs in the combined group of NDD (SCZ, ID/DD and ASD). This was especially true when restricting to the LoF-intolerant genes (pLI > 0.9 or LoF-FDR < 0.01) hit by LoF DNMs (Figure 4B, *P* = 1.0 × 10^-6^, FC = 2.1 with pLI, *P* < 1.0 × 10^-6^, FC = 2.1 with LoF-FDR). Significant enrichment was not observed among the genes hit by LoF DNMs, irrespective of their selective pressure, in unaffected subjects (Figure 4B, smallest *P* = 0.42) or congenital heart disease patients (Figure 4B, smallest *P* = 0.24). Stratification of NDD by disease, revealed an enrichment in ID/DD, ASD and SCZ with variable significance probably due to the number of input DNMs and the consequent statistical power (Figure 4B).

We further searched for disease-related functional and expression properties of Setd1a target genes within the conserved and chromatin accessible genomic regions used in our analysis. GO analysis of high-confidence Setd1a target genes revealed a significant enrichment in terms relevant to “ion channel activity”, “neurogenesis”, “synaptic function”, as well as significant enrichment for the “schizophrenia” term as defined in the DisGeNET database (umls:C0036341) (Figure 4C). Brain regions and developmental stage analysis using the CSEA tool (see Methods) showed a significant enrichment primarily in the cortex at early fetal/ early mid fetal stages (FDR corrected pSI *P* = 0003 and *P* = 0.002, respectively) and to a lesser extent at the young adulthood stage (pSI *P* < 0.05) (Figures 4D). Notably removal of Setd1a target genes overlapping with genes carrying LoF DNMs in NDDs, which are prenatally-biased in their expression (Fig S5E), resulted in stronger enrichment at young adulthood/adolescence stages (FDR corrected pSI *P* = 2.544e-07 and 7.600e-06, respectively) (Figures 4E), suggesting that Setd1a targets genes with distinct roles in both prenatal and postnatal neurodevelopment.

### Setd1a targets display a complex pattern of transcriptional dysregulation

To obtain insights into the molecular alterations in the PFC of *Setd1a^+/-^* mice, we used RNA-Seq analysis to profile transcriptomes of micro-dissected PFCs from four 6-week-old *Setd1a^+/-^* mice and four WT littermates. Changes in gene expression in complex adult tissues, consisting of heterogeneous cellular populations, represent a composite state of primary effects on gene expression in combination with secondary compensatory and downstream transcriptional events (i.e. due changes in local activity) as well as with alterations due to earlier developmental insults. We identified 342 genes that significantly change their expression in *Setd1a^+/-^* mice (FDR<5%) (Figure 5A, Data Table 1). Transcriptional alterations were modest with most of the genes (> 70%) displaying a fold change in the range of 0.7-1.3. Interestingly, the majority of the differentially expressed genes (79%, 271/342) in *Setd1a^+/-^* PFC overlap with direct targets of Setd1a, likely representing primary rather than secondary transcriptional alterations.

**Figure 5.**
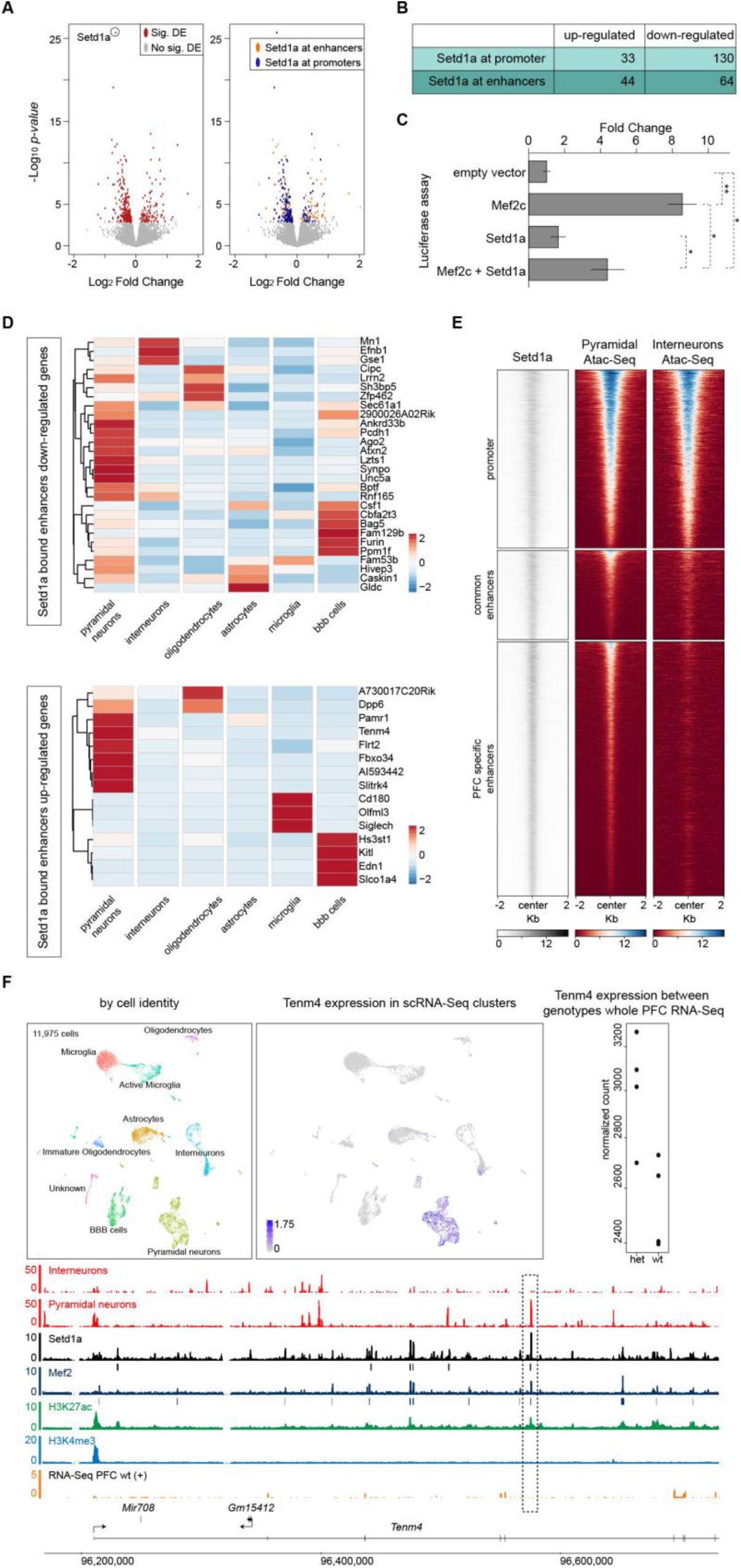
Characterization of differentially expressed genes in PFC of *Setd1a^+/-^* mice and cell-type specific expression of Setd1a target genes. (A-B) Characterization of differentially expressed genes in PFC of *Setd1a^+/-^* mice. Volcano plots of differentially expressed genes in whole-PFC RNA-Seq of *Setd1a^+/-^* mice are depicted in (A). 342 differentially expressed genes (FDR<5%) are shown in red, while non-significant genes are shown in grey (*left panel*). Of the 342 differentially expressed genes, 271 are Setd1a targets (orange and blue, *right panel*). We partitioned these genes in 4 categories based on the binding regions of Setd1a in the PFC. We defined ‘Setd1a bound promoters up-regulated genes’ and ‘Setd1a bound promoters down-regulated genes’ as genes in which promoters are bound by Setd1a. We defined ‘Setd1a bound enhancers up-regulated genes’ and ‘Setd1a bound enhancers down-regulated genes’ as genes in which enhancers, but not promoters, are bound by Setd1a. To avoid counting replicates, genes with Setd1a binding at both promoters and enhancers, were partitioned into the “Setd1a-promoter bound” classes. A summary of the number of genes in each category is shown in (B). (C) Luciferase reporter assay of Mef2 responding element when Mef2c and/or Setd1a are co-transfected transiently in HEK293T cells. While Mef2c alone showed 8.55 fold increase activity over the background, when Mef2c is co-transfected with Setd1a, we observed a significant decreased activity to 4.47 fold. Values are mean ± standard error for three biological replicates, normalized to the empty vector (see methods). Student’s t-test, *P < 0.05, **P < 0.01. (D) Heatmaps of the expression levels of down-regulated (*top panel*) and up-regulated (*bottom panel*) genes with Setd1a bound enhancers, across different cell types identified by scRNA-Seq (see methods and Figure S6C and S6D). For this analysis we combined the two different types of microglia and oligodendrocytes that we identified. Heatmaps show the normalized UMI counts per genes of the two biological replicates per genotype (see Methods). (E) Heatmap of Setd1a ChIP-Seq coverage and ATAC-Seq of sorted pyramidal neurons and interneurons, 2 kilobases upstream and downstream of the center of Setd1a bound promoters (*top panel*) and enhancers (*bottom panel*). All heatmaps are sorted in the same order, based on the Setd1a signal. (F) *Tenm4* locus is shown as an example of genes that are up-regulated in Setd1a mutant PFC and are selectively expressed in pyramidal neurons. UMAP representation of 11,975 single cell transcriptomes from the PFC of WT (5,944) and *Setd1a^+/-^* (6,031) mice, colored by the identified cell-types (see Methods, Figure S5C and S5D), is shown on the *top-left panel*. Tenm4 is specifically expressed in the pyramidal neuron scRNA-Seq cell cluster (*top-middle panel*), and whole PFC RNA-Seq showed that *Tenm4* is up-regulated in Setd1a mutant mice (*top-right panel;* each dot is the normalized expression of *Tenm4* in each biological replicate per genotype). Representative RNA-Seq data of one biological replicate of WT mice (*orange*), ChIP-Seq of Mef2 (*dark blue*), Setd1a (*black*), H3K27ac (*green*), H3K4me3 (*light blue*) as well as ATAC-Seq from pyramidal neurons and interneurons (*light red*) are shown (*bottom panel*). Black and dark blue boxes, below Setd1a and Mef2 tracks respectively, represent significant peaks that passed all quality checks (see Methods). Highlighted with a dashed square is the regulatory element that is bound by Setd1a and Mef2, marked by H3K27ac but not by H3K4me3, and accessible in pyramidal neurons but not in interneurons.

To dissect the effect of the *Setd1a* mutation on gene expression, we partitioned differentially expressed genes based on the binding regions of Setd1a in the PFC. Genes were classified as up- or down-regulated and carrying Setd1a binding sites at promoters or enhancers (Figure 5A and 5B). As anticipated from the well-known function of Setd1a as transcriptional activator (Shilatifard, 2012), the majority (78%) of genes with Setd1a-bound promoters were down-regulated in *Setd1a^+/-^* mice (Figure 5B). In contrast, genes that have Setd1a-bound intergenic enhancers can be either up-(40%) or down-regulated (60%) (Figure 5B). Although the complex pattern of transcriptional changes in the PFC of Setd1a^+/-^ mice could be explained by a combination of direct and indirect effects, we explored the possibility that Setd1a may have opposing functions and molecular targets in promoters and enhancers. Because Setd1a binding shows an almost complete overlap with Mef2 binding on enhancer sequences, and because Mef2 activity can be inhibited by lysine methylation (Choi et al., 2014), we reasoned that Setd1a may act likewise as a Mef2 repressor on enhancer elements restricting its transactivation activity (while it acts as activator on promoters via H3K4 trimethylation). Consistent with this hypothesis, Setd1a inhibits Mef2c-mediated transcriptional activation in transient transfection assays (that remove the influence of histone methylation, Figure 5C), providing a possible explanation for the observed transcriptional up-regulation of genes with Setd1a-bound enhancers upon deletion of one Setd1a allele.

In addition to the presumed opposing functions of Setd1a on promoters and Mef2-bound enhancers, cellular heterogeneity of the PFC may also contribute to the modest and heterogeneous transcriptional effects observed at the PFC of *Setd1a*^+/-^ mice. To test this, we performed single cell RNA-Seq (scRNA-Seq) on dissociated cells from the PFC of WT and *Setd1a^+/-^* mice. We obtained expression profiles for 5,944 cells from WT mice and 6,031 cells from *Setd1a^+/-^* mice (with a mean of 2262 genes per cells). We identified 8 main clusters that represent the major neuronal [pyramidal neurons (*Camk2a*) and interneurons (*Igfbpl1*)] and non-neuronal [mature and immature oligodendrocytes (*Olig2* and *Mbp*), microglia (*Aif1*), astrocytes (*Aldoc*) and blood brain barrier cells (*Cldn5*)] populations of PFC (Figures S6A, S6B, S6C and S6D). As an indicator for the quality of our scRNA-Seq data, we compared expression, as a whole, of genes that we detected as significantly changing in bulk PFC RNA-Seq and confirmed the same trends of expression changes between bulk and scRNA-Seq (Figures S6Eand S6F). Differential expression analysis failed to detect robust cell type-specific genotypic changes in the expression of genes proximal to Setd1a peaks and did not reveal significant changes in the expression of individual genes. While this finding may be in part due to well-known limitations in sensitivity and technical noise of scRNA-Seq that may mask underlying biological variation (Liu and Trapnell, 2016), it strongly suggests that reduction of Setd1a levels by half leads to low magnitude changes in the expression of downstream targets even at the cellular level.

We used the scRNA-Seq data to dissect the cellular origins of the expression alterations identified by bulk RNA-Seq and highlight potential targets underlying the phenotypic alterations we observed in cortical pyramidal neurons. Analysis of cellular distribution of all up- and down-regulated genes in *Setd1a^+/-^* mice revealed a heterogeneous peak expression pattern spanning several neuronal as well as pathophysiologically relevant (Chen et al., 2018; Volk, 2017; Windrem et al., 2017) non-neuronal cell-types, such as oligodendrocytes, astrocytes and microglia (Figure 5C, S6G and S6H). However, pyramidal cells display the highest fraction of differentially expressed genes compared to any other cell type identified by scRNA-Seq, consistent with the observation that genes that have Setd1a peaks at their intergenic enhancers have their highest expression in pyramidal neurons (Figure S6I). The transcriptional dysregulation of some of these pyramidal neuron-specific and psychiatric disease relevant genes, such as Slitrk4 (Proenca et al., 2011) (an up-regulated gene involved in the repression of neurite growth) as well as *Synpo* (Kos et al., 2016) and Unc5a (Rajasekharan and Kennedy, 2009) (downregulated genes involved in synapse and dendritic spine formation and maintenance) (Figure 5D) could contribute to the morphological and synaptic alterations observed in *Setd1a^+/-^* mutant mice.

The correlation between pyramidal neuron-specific expression and proximity to Mef2/Setd1a-bound intergenic peaks immediately suggests that many of these genomic regions might represent pyramidal-neuron specific regulatory elements. To test this, we FAC-sorted pyramidal neurons and interneurons from double transgenic mice (from either Emx1-ires-Cre or Parvalbumin-ires-Cre lines crossed to tdTomato reporter mice) and analyzed them by ATAC-Seq. This approach confirmed that Sedt1a-bound intergenic peaks are accessible and occupied by transcription factors specifically at pyramidal neurons (Figures 5E). As an example, the *Tnem4* gene locus is shown in more detail in Figure 5F. This gene is expressed in PFC pyramidal neurons and has been found to be up-regulated in *Setd1a*^+/-^ mice, with Setd1a bound at its enhancers but not at its promoter. We identified one enhancer element (regions marked by H3K27ac but not H3K4me3) that is bound by both Setd1a and Mef2c specifically in pyramidal neurons and not in interneurons, as shown by our cell-type specific ATAC-Seq.

### Reinstating *Setd1a* expression in adulthood rescues WM deficits

The lack of evidence of neurodegenerative changes or gross structural abnormalities in *Setd1a*-deficient brains prompted us to investigate the possibility of reversing cognitive deficits in adulthood. To this end, we took advantage of the conditional nature of the targeted allele and adopted a genetic method that allows for inducible reactivation of Setd1a in *Setd1a^+/−^* adult mice (see Methods). To achieve temporal control, we crossed a *Setd1a^+/-^* mice to an inducible *R26^FlpoER^* mouse line (*R26FlpoER*) that activates global Flpo function upon tamoxifen treatment (Figures 6A). The *Flp/Frt*-dependent genetic switch strategy, which enables the conditional manipulation of *Setd1a* at its endogenous genomic locus, thus preserving *Setd1a* expression within physiological levels. When the *Setd1a^+/-^: R26FlpoER^+/-^* mice reached 8 weeks, we used oral gavage to deliver tamoxifen and implement Flpo-mediated gene restoration (Figures 6A and 6B).

**Figure 6.**
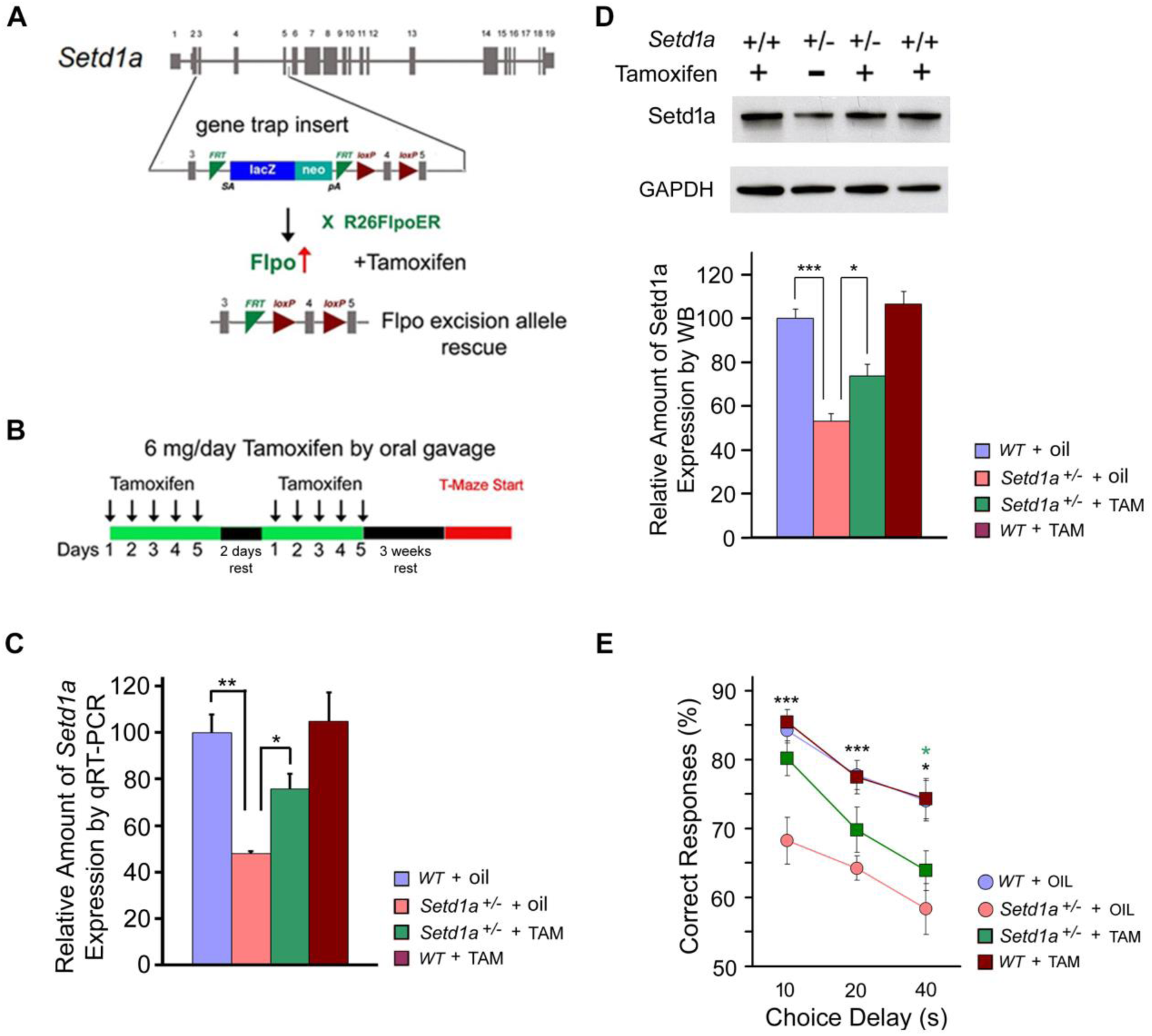
Restoration of Setd1a expression in adulthood rescues WM performance. (A) Schematic of the restoration of knockout-first allele after tamoxifen treatment. For the rescue experiments, *Setd1a^+/-^* mice were crossed to *R26FlpoER* mice. FLP (Flpo) induced globally by tamoxifen converts the “knockout-first” allele to a conditional allele, restoring gene activity. (B) Tamoxifen feeding scheme. Animals were fed tamoxifen for 5 consecutive days followed by 2 days of rest, then fed for 5 more days followed by 3 weeks of rest before the T-maze assay. (C) qRT-PCR of *Setd1a* expression in PFC of adult mice, demonstrating the restoration of *Setd1a* mRNA levels following tamoxifen treatment. N = 6 mice each genotype, n = 3 each animal. Data are shown as means ± s.e.m. \**P* < 0.05; \*\**P* < 0.01. Student’s two-tailed *t*-test. (D) Western blot analysis of Setd1a levels in lysates from PFC of adult mice, demonstrating the restoration of Setd1a protein levels by tamoxifen treatment. N = 4 mice each genotype, n = 3 each animal. Data are shown as means ± s.e.m. \**P* < 0.05; \*\**P* < 0.01, \*\*\**P* < 0.001. Student’s two-tailed *t*-test. (E) WM performance during testing phase. Mean percentage of correct responses for WT and *Setd1a^+/^* mice with/without tamoxifen treatment. *\*P*, 0.05, *\*\*P*, 0.01 (black asterisk: WT + oil versus *Setd1a^+/-^* + oil, green asterisk: *Setd1a^+/-^* + oil versus *Setd1a^+/-^* + TAM).

Behavioral analysis was performed on the following 4 groups: *Setd1a*^+/+^: *R26FlpoER*^+/−^ mice treated with tamoxifen (WT + TAM), *Setd1a*^+/-^:*R26FlpoER*^+/−^ mice treated with tamoxifen (*Setd1a*^+/-^ + TAM), *Setd1a*^+/+^:*R26FlpoER*^+/−^ mice treated with corn oil vehicle (WT + oil), and *Setd1a*^+/-^:*R26FlpoER*^+/−^ mice treated with vehicle (*Setd1a*^+/-^+ corn oil). Investigation of the efficiency of Setd1a reactivation upon Flpo-mediated deletion of the *LacZ* cassette confirmed that upon tamoxifen treatment Setd1a mRNA and protein levels were reinstated in *Setd1a*^+/-^ + TAM mice to ∼75% of WT levels in the adult mouse brain (Figures 6C, 6D, S7A, S7B, and S7C). The effect of elevation of Setd1a levels on spatial WM performance was assessed in a T-maze delayed non-match to place task. As expected, *Setd1a*^+/-^+ oil mice showed impaired WM performance compared to WT + oil control littermates, which was partially rescued upon tamoxifen treatment (Figures 6E, S7D, and S7E). Rescue was particularly evident at 10s in *Setd1a*^+/-^ + TAM mice (82.64 ± 2.29%, N = 24) compared to WT + TAM littermates (85.42 ± 1.83%, N = 24), or *Setd1a*^+/-^+ oil (68.25 ± 3.38%, N = 21) (*P* = 0.35 versus WT + TAM; *P* = 0.0012 versus *Setd1a*^+/-^ + oil), but only modestly (statistically non-significant) at longer delays.

Partial restoration of neurocognitive deficits might have resulted from the incomplete reinstatement of Setd1a expression (∼75% of WT levels) or could reflect a partial developmental contribution requiring additional earlier interventions. These limitations notwithstanding our findings indicate that, consistent with continued plasticity in the adult brain (Mei et al., 2016), adult *Setd1a* function reinstatement has substantial effect on ameliorating neurocognitive deficits and highlights a broad window of therapeutic opportunity (Neale et al., 2012). We explored this possibility further by taking advantage of the more homogeneous biodistribution of systemically administered pharmacological compounds that overcome the limited efficiency of genetic manipulations.

### Pharmacological Inhibition of LSD1 demethylase activity in adulthood counteracts the effects of *Setd1a* deficiency

Within the past few years, a number of histone demethylases have been discovered, as well as an ever-increasing number of potent and selective histone demethylases inhibitors that exhibit good brain penetration. We thus surmised that these compounds would most likely generate robust methylation changes, which could, in turn, counteract effects of *Setd1a* deficiency and lead to restoration of Setd1a-linked deficits. Setd1a-counteracting demethylase(s) have yet to be described. We hypothesized that LSD1 may be one such demethylase since both our whole tissue RNA-Seq and scRNA-Seq analysis identified LSD1 as the most abundant demethylase in the frontal cortex. To test whether LSD1 binds *in vivo* the same regulatory elements bound by Setd1a we performed LSD1 ChIP-Seq from the prefrontal cortex of 6 weeks old mice, identifying 43,917 LSD1 bound regions (IDR<0.05). Strikingly, as for Mef2, we observed a compelling overlap between LSD1 and Setd1a bound regions (22,532 out of 25,658 are co-bound by LSD1), both at promoter and enhancer elements (Figures 7A, 7B, and 7C, the *Neurod6* locus is shown as an example in Figure 7A). Notably, in agreement with previous observations that Mef2 activity can be inhibited by lysine methylation (Choi et al., 2014), we showed that LSD1 enhances Mef2c-mediated transcriptional activation in transient transfection assays (Figure 7D), an effect opposite to the one observed for Setd1a (Figure 5C), which supports the view that Setd1a and LSD1 can modulate Mef2 activity in opposite directions independently of their effects on histones.

**Figure 7.**
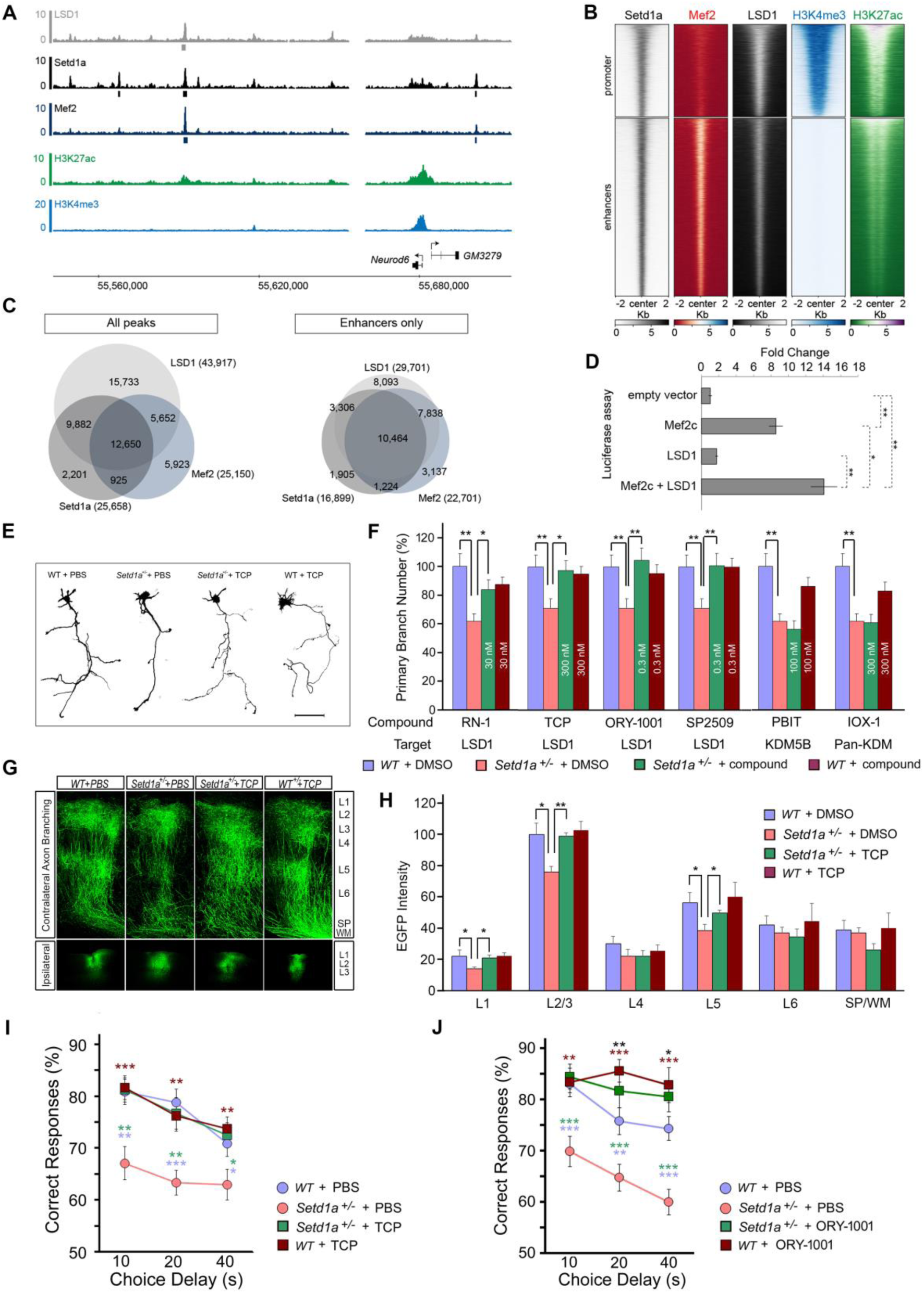
Inhibition of LSD1 activity in adulthood counteracts the effects of *Setd1a* deficiency. (A) *Neurod6* locus. ChIP-Seq of LSD1 (*grey*), Setd1a (*black*), Mef2 (*dark blue*), H3K27ac (*green*), H3K4me3 (*light blue*) is shown. Grey, black and dark blue boxes below Lsd1, Setd1a and Mef2 tracks respectively represent significant peaks that passed all quality checks (see methods). (B) Heatmap of the ChIP-Seq coverage for Setd1a, LSD1, Mef2, H3K4me3 and H3K27ac 2 kilobases upstream and downstream of the center of Setd1a peaks [promoters (*top panel*) and enhancers (*bottom panel*)]. All heatmaps are sorted in the same order, based on Setd1a and H3K27ac signals. (C) Venn diagram of the overlap between all the peaks (*left panel*) and enhancers only peaks for LSD1, Setd1a and Mef2 ChIP-seq (*right panel*). (D) Luciferase reporter assay of the Mef2 responding element when Mef2c and/or LSD1 are co-transfected in HEK293T cells. While Mef2c alone showed 8.55 fold increase activity over the background, when Mef2c is co-transfected with LSD1, we observed a significant increased activity to 14 fold. Values are mean ± standard error for three biological replicates, normalized to the empthy vector (see methods). Student’s t-test, *P < 0.05, **P < 0.01. (E and F) *Setd1a^+/-^* cortical neurons were treated with LSD1/KDM1a, or JmjC domain KDM inhibitors from DIV3 to DIV5, and transfected with plasmids expressing EGFP at DIV4 and fixed at DIV5. Tracings of representative neurons imaged at DIV5 following TCP or PBS treatment are depicted in (E). The most effective concentrations of each inhibitor for restoring primary axon branch number in *Setd1a^+/-^* neurons (0.1 nM – 30 μM) are represented in (F). There are no significant differences between WT + DMSO and WT + compound in any of the comparisons.*\*P*, 0.05, *\*\*P*, 0.01. Histograms show means and error bars represent s.e.m. Scale bar, 50 μm. (G and H) TCP treatment restores axon branch number *in vivo*. (G) Representative images of contralateral axon-terminal branching of EGFP-labeled neurons from coronal sections of adult brains of mice injected AAV2Synapsine-EGFP virus at 8 weeks (*upper panels*) and viral injection sites in ipsilateral L2/3 (*bottom panels*). Immuno-histological staining represents EGFP (*green*). (H) Histograms depicting quantitative assessment of contralateral axon branching in L1 to L6 and white matter in adulthood and indicating that TCP treatment restores contralateral axon branching in L2 to L5 in *Setd1a^+/-^* mice. WT + PBS: N = 6, *Setd1a^+/-^* + PBS: N = 7, *Setd1a^+/-^* + TCP: N = 6, WT + TCP: N = 5. *\*P*, 0.05, *\*\*P*, 0.01. Histograms show means and error bars represent s.e.m. (I) Complete rescue of WM performance in a delayed non-match to place task in *Setd1a^+/-^* mice treated with TCP. Mean percentage of correct responses for WT and *Setd1a^+/-^* mice with/without 3 mg/kg/day TCP or vehicle treatment are depicted. *\*P*, 0.05. *\*\*P*, 0.01, *\*\*\*P*, 0.001 (*blue asterisk:* WT + PBS *vs Setd1a^+/-^* + PBS, *green asterisk: Setd1a^+/-^* + TCP *vs Setd1a^+/-^* + PBS*, brown asterisk:* WT + TCP *vs Setd1a^+/-^* + PBS). No significant difference between WT + PBS and *Setd1a^+/-^* + TCP groups was observed. Data are mean ± s.e.m (*Setd1a^+/-^* + PBS: N = 20, *Setd1a^+/-^* + TCP: N = 20, WT + PBS: N = 20, WT + TCP: N = 20). (J) Rescue of WM performance in a delayed non-match to place task in *Setd1a^+/-^* mice treated with ORY-1001. Mean percentage of correct responses for WT and *Setd1a^+/-^* mice with/without 10 μg/kg/day ORY-1001 or vehicle treatment are depicted. *\*P*, 0.05, *\*\*P*, 0.01, *\*\*\*P*, 0.001 (*blue asterisk:* WT + PBS *vs Setd1a^+/-^* + PBS, *green asterisk: Setd1a^+/-^* + ORY-1001 *vs Setd1a^+/-^* + PBS*, brown asterisk:* WT + ORY-1001 *vs Setd1a^+/-^* + PBS, *black asterisk:* WT + ORY-1001 *vs* WT + PBS). No significant difference between WT + PBS and *Setd1a^+/-^* + ORY-1001 groups was observed. Data are mean ± s.e.m (*Setd1a^+/-^* + PBS: N = 21, *Setd1a^+/-^* + ORY-1001: N = 15, WT + PBS: N = 23, WT + ORY-1001: N = 15).

To confirm the ChIP-Seq findings we undertook parallel phenotypic screens on primary cultures of *Setd1a^+/-^* cortical neurons using well-characterized demethylase inhibitors and determining their potential to rescue the robust axonal arborization deficit (Figures 7E and 7F). *Setd1a^+/-^* neurons were treated with inhibitors of LSD1/KDM1a [RN-1, tranylcypromine (TCP), ORY-1001, and SP2509], KDM5B (PBIT) and the pan-KDM inhibitor IOX-1 from DIV3 to DIV5, transfected with plasmids expressing EGFP at DIV4 and fixed at DIV5. TCP, ORY-1001, SP2509, and RN-1 (all LSD1 inhibitors), but not PBIT or IOX-1 rescued the axon branching deficits in *Setd1a^+/-^* neurons, confirming an important role of LSD1 antagonism in counteracting the effects of *Setd1a* deficiency (Figures 7E and 7F).

We followed up the LSD1 findings by examining the effect of inhibition of LSD1 on cognitive deficits following systemic TCP administration (Figures 7I, S7F, S7G, S7H, S7I, and S7J). *Setd1a^+/−^* mice and WT littermates were injected with 3 mg/kg/day TCP or vehicle for 14 consecutive days followed by the non-match to sample T-maze task. During the task, mice were administered daily with the same dose of TCP or vehicle. Assessment of spatial WM performance revealed modest genotypic effects during training (Figures S7F and S7G), but a robust impairment in the performance of vehicle-treated mutant mice over vehicle treated WT control littermates during testing with increasing time delays (Figure 7I). This deficit was completely reversed by TCP treatment, such that TCP-treated *Setd1a^+/−^* mice performed indistinguishably from TCP-treated WT littermates (Fig. 7I). TCP had no effect on WT mice. While vehicle treated mutant mice showed the expected deficits in days to criterion and correct percentages during training phase, TCP treatment did not improve these deficits. Follow-up dose-response studies indicated that the effective TCP dose range was rather narrow: a lower dose (1 mg/kg/day, i.p.) was ineffective in improving the WM performance (Figures S7H, S7I and S7J) whereas a higher dose of TCP (10 mg/kg/day, i.p.) resulted in occasional lethality precluding further cognitive assessment.

Pharmacological inhibition of LSD1 *in vitro* can repair axonal terminal branching but it is unclear whether *in vivo* post-developmental LSD1 antagonism by TCP has similar beneficial effects on the attenuated terminal arbor complexity, parallel to the rescue of the cognitive deficits. To address this question, we injected adeno-associated virus rAAV2/hSynapsin/EGFP into the frontal cortex of 12-14-week-old mice following the T-maze test to label cortical neurons with EGFP in TCP- or vehicle-treated mutant and WT mice. Mice were sacrificed after 2 weeks incubation. We analyzed EGFP intensity in contralateral axon target regions as a quantitative index of the complexity of terminal axon branching. As expected mutant mice treated with vehicle showed significantly reduced callosal axon-terminal branching within the contralateral cortical L2/3 (Figures 7G and 7H). By contrast TCP treatment restored axon branching in contralateral L2/3 of *Setd1a^+/-^* to near WT level (Figures 7G and 7H).

While TCP is a prototype irreversible LSD1 inhibitor, it also inhibits two major isoforms of monoamine oxidases MAO-A and MAO-B. To eliminate possible confounding effects of MAO inhibition on WM we also evaluated the effect of ORY-1001, a TCP-derivative that represents the most potent and selective LSD1 inhibitor described to date (Hosseini and Minucci, 2017; Maes et al., 2015; Maes et al., 2018). The compound is an irreversible inhibitor and has an IC50 of 18 nM and an over 1000-fold selectivity for LSD1 compared to MAOs (Maes et al., 2015; Maes et al., 2018). Indeed, in our in vitro axon branching rescue assays ORY-1001 was 1000-fold more potent than TCP and demonstrated sub-nanomolar cellular efficacy (Figure 7F). To examine whether ORY-1001 is effective in reversing WM deficits in *Setd1a^+/−^* mice, mutant mice and WT littermates were administered with ORY-1001, or vehicle (PBS) for 14 consecutive days prior to the T-maze task and daily throughout the duration of the task. As expected, we observed a robust impairment in the performance of vehicle-treated mutant mice over vehicle treated WT control littermates during testing with increasing time delays. This WM deficit was reversed by 10 μg/kg/day ORY-1001 treatment of *Setd1a^+/−^* mice (Figure 7J). Days to criterion and percentage of correct choices during training phase were also improved in both ORY-1001-treated *Setd1a^+/−^* and WT mice (Figures S7K and S7L). Notably, administration of lower of ORY-1001 (3 μg/kg/day) proved ineffective in improving the WM performance during testing with increasing time delays (Figures S7M, S7N, and S7O).

## Discussion

The discovery of rare but highly penetrant genetic risk variants for SCZ and the generation of mouse models that faithfully recapitulate their effects has provided unprecedented insights into disease pathophysiology (Arguello and Gogos, 2012; Del Pino et al., 2018). A major remaining challenge is how to begin to translate these mechanistic insights into effective therapies. In that context, this study describes a mouse model of *SETD1A* LoF SCZ-risk mutations and evaluates *how* a bona fide pathogenic mutation, which disturbs a relatively ubiquitous molecular process, affects the structure and function of cortical neural circuits and *whether* these effects are reversible in the adult brain.

We describe a profile of robust but circumscribed cognitive impairments in spatial WM as well as cortical synaptic alterations due to *Setd1a-*happloinsufficiency, including sparser structural connectivity due to reduction in terminal arborization of cortical axons as well as alterations in short-term synaptic plasticity and neuronal excitability, which collectively may lead to disruptions of network activities that support WM and related cognitive operations (Mi et al., 2017). Overall, *Setd1a^+/-^* phenotypes relate well to established evidence that SCZ patients have reduced WM capacity (Arguello and Gogos, 2012) as well as to cognitive and synaptic alterations described in two other models of highly penetrant SCZ-risk mutations (Crabtree et al., 2017; Fenelon et al., 2013; Koike et al., 2006; Mukai et al., 2008; Stark et al., 2008). Comparison among these strains, together with previous *in vivo* analysis of neuronal activity of locally (Hamm et al., 2017) or distally (Tamura et al., 2016) connected neocortical populations, suggests a pattern of dysconnectivity within and between brain regions, shaped by alterations in rapid synaptic dynamics, altered intrinsic excitability and sparser connectivity, which may affect the coherence of neuronal activity and may underlie the emergence of cognitive deficits (Gogos et al., 2019).

Our ChIP-Seq analysis, revealed a large number of putative Setd1a direct chromatin targets, distributed over enhancer and promoter elements. Setd1a-bound enhancers, in particular, appear to have a cortex- and neuronal-specific function both in mouse and human, suggesting that the molecular mechanisms that lead to the recruitment of Setd1a to enhancers are conserved between species. This finding, as well as data from chromatin accessibility (ATAC-Seq) assays indicating that many Sedt1a enhancer peaks are accessible specifically in pyramidal neurons, strongly imply the presence of auxiliary factors that may mediate the brain region and cell-type-specific effects of this relatively ubiquitously expressed regulator. Indeed, our ChIP-Seq and immunoprecipitation analysis revealed a key role for Mef2 transcription factors in mediating the transcriptional effects of Setd1a enhancer occupancy. Notably, Mef2 isoforms have been shown to modulate cortical structural and functional plasticity and have been previously implicated in SCZ, ASD and other neuropsychiatric disorders (Harrington et al., 2016; Mitchell et al., 2017) inviting to speculate that Setd1a and Mef2 may be working in concert to regulate expression of disease-relevant genes. Interestingly, genes with Setd1a-bound enhancers show a highly specific overlap with established lists of psychiatric and neurodevelopmental disease risk genes suggesting that *Setd1a* haploinsufficiency in both mice and humans may lead to behavioral and physiological deficits in part via coordinated mis-regulation of disease-risk genes.

Interpreted in the context of our bulk and scRNA-Seq analysis of PFC, reduction of Setd1a levels by half may lead to low magnitude changes in the expression of many downstream targets with likely cumulative effects on cellular structure and function, especially in pyramidal neurons where Setd1a enhancer-bound targets show the highest levels of expression. Given the previously described role of the COMPASS complex as transcriptional activator, one unexpected finding from our RNA-seq expression analysis, is that reduction of Sedt1a levels by half results to a large extent in upregulation of direct target genes in addition to predicted downregulation. Specifically, while genes with Setd1a-bound promoters were predominantly down-regulated in *Setd1a^+/-^* mice, a large fraction of genes that have Setd1a-bound enhancers were up-regulated. While this finding requires further investigation, it is possible that Setd1a may have other functions as a methyl-transferase, in addition to catalyzing methylation of H3K4. For example, recent data suggest that Mef2 can be regulated by histone methyltransferase G9a-dependent lysine methylation, which inhibits its activity (Choi et al., 2014; Ow et al., 2016). In that vein, it is possible that Setd1a also methylates Mef2, negatively regulating its activity in some Mef2/Setd1a-cobound enhancers. Consistent with this interpretation we provide evidence that Setd1a inhibits Mef2c-mediated transcriptional activation in transient transfection assays, providing a possible explanation for the observed transcriptional upregulation of genes with Setd1a-bound enhancers upon deletion of one Setd1a allele.

Notably, while some of the alterations in *Setd1a^+/-^* mice may be caused by impaired prenatal development, one of the novel aspects of the current study was the demonstration that specific cognitive and neural circuitry impairments in the Setd1a-deficient brain can be partially or fully overcome by genetic or pharmacological interventions aiming at delayed restoration of Setd1a function. Our results revealed a beneficial effect of post-developmental activation of *Setd1a* expression on cognitive dysfunction, highlighting a unique role of Setd1a expression after development and arguing that the deficiency of Setd1a function during early developmental stages does not inexorably lead to irreversible cellular and behavioral alterations. Similar analysis in other models of monogenic brain disorders has provided mixed results. For example, adult reactivation of *Ube3a* or the *Syngap1* gene in mouse models of autism and intellectual disability did not reverse any of the core behavioral deficits (Barnes et al., 2015; Rotaru et al., 2018), while adult reactivation of *Mecp2* could rescue behavioral alterations in a mouse model of Rett syndrome (Tillotson et al., 2017). Guided by these findings, we employed a series of effective pharmacological interventions to directly target, with well-characterized and potent inhibitors, downstream demethylation processes. Our goal was to generate robust methylation changes which, in turn, could counteract effects of *Setd1a* deficiency leading to restoration of Setd1a-linked deficits of cell function and perhaps behavior. Combined results from ChIP-Seq analysis and *in vitro* phenotypic screens identified LSD1 as a major demethylase counteracting the effects of Setd1a methyl transferase activity. Further *in vivo* analysis showed that LSD1 antagonism in *Setd1a*-deficient mice results in a full rescue of the behavioral abnormalities and axonal arborization deficits. While it will be important to determine the specific methylation sites underlying the beneficial effects of LSD1 antagonism, our findings indicate that reactivating Setd1a function or counteracting downstream effects of *Setd1a* deficiency in the adult brain may be sufficient to ameliorate the manifestation of neurocognitive phenotypes highlighting the continued structural and functional plasticity in the adult brain. Thus, in addition to mechanistic insights, our findings may prove useful for development of new therapeutic strategies. Therapies targeted to SETD1A may have wider implications as suggested by our findings here as well by the fact that pathways related to histone methylation are strongly enriched among biological pathways in GWAS data of SCZ (Network and Pathway Analysis Subgroup of Psychiatric Genomics, 2015).

## Author Contributions

J.M., E.C., B.X., A.T., S.L., and J.A.G designed and interpreted the experiments. J.M validated the mouse model, performed, analyzed and interpreted behavioral, morphological, co-IP, targeted expression and all rescue assays. E.C. performed, analyzed and interpreted all ChIP-Seq and scRNA-Seq assays. G.W.C. and Z.S. performed, analyzed and interpreted electrophysiology experiments. A.D performed, analyzed and interpreted behavior experiments. P.T contributed to the *in vivo* axonal branching assays. C-Y.C performed and analyzed targeted expression assays. B.X performed the bioinformatics analysis of the overlap between ChIP-Seq peaks and risk genes. A.T analyzed the RNA-Seq data and contributed to the validation of the mouse model and the bioinformatics analysis of the ChIP-Seq data. Y.C contributed the bioinformatics analysis. S.L. and J.A.G supervised the work. J.M., E.C., G.W.C., Z.S., A.D, A.T., B.X., S.L., J.A.G wrote the paper. The authors declare no competing financial interests.

## Acknowledgements

We thank Dr. Sneha Rao for her help with in vivo axon imaging and Naoko Haremaki for the maintenance of the mouse colony and technical assistance. This work was partially supported by grants R01MH080234 (to J.A.G), R01DA036894 (to S.L).

## STAR Methods

### LEAD CONTACT AND MATERIALS AVAILABILITY

Information and requests for resources and reagents should be directed to, and will be fulfilled without restriction by, the Lead Contact Joseph A. Gogos.

### EXPERIMENTAL MODEL AND SUBJECT DETAILS

*Setd1a^tm1a(EUCOMM)Wtsi^* mice (referred to as *Setd1a^+/-^*) were obtained from EMMA (https://www.infrafrontier.eu/search). *Setd1a^+/-^* mice were backcrossed for 10-11 generations in the C57BL/6J (The Jackson Laboratory, Bar Harbor, ME) background. B6N. 129S6(Cg)-*Gt(ROSA)26Sor^tm3(CAG-flpo/ERT2)Alj^*/J (referred to as *R26^FlpoER^)*, Emx1tm1(cre)Krj/J, Palvtm1(cre)Arbr/J, and Gt(ROSA)26Sortm14(CAG-tdTomato)Hze/J were purchased from Jackson laboratory. To achieve temporal control of *Setd1a* expression for the *Setd1a* rescue experiments, we crossed the *Setd1a^+/^* mice to an inducible *R26^FlpoER+/+^* (Lao et al., 2012) mouse line that activates global *Flp* function after tamoxifen treatment. B6.Cg-*Gt(ROSA)^26Sortm1.3(CAG-tdTomato,-EGFP)Pjen^*/J (referred to as *RC::FLTG*) (Plummer et al., 2015) was purchased from Jackson Labs. To evaluate the tamoxifen-inducible Flp recombination in neurons, we crossed the *RC::FLTG* mice to an inducible *R26^FlpoER+/+^* (Lao et al., 2012) mouse line that activates global *Flp* function after tamoxifen treatment. For ATAC-Seq experiment Emx-Cre mice (Jackson lab) and Parv-Cre mice (Jackson lab) were crossed to CRE inducible tdTomato mice (Jackson lab) so that pyramidal neurons or parvalbumin expression interneurons expressed the fluorescent protein tdTomato

Animals were housed in groups of five per cage, under a 12-h light/12-h dark cycle, with controlled room temperature and humidity, and then housed singly for behavioral experiments. Food and water were provided *ad libitum*. All behavioral experiments were performed on adult male mice aged 8–14 weeks during the light cycle. All animal procedures were carried out in accordance with and approved by the Columbia University Institutional Animal Care and Use Committee.

### METHOD DETAILS

#### Co-immunoprecipitation

Nuclear extract was prepared from fresh cortex of 8-wk old mice using Nuclear Complex Co-IP Kit (Active Motif. Inc.) according to the manufacturer’s instructions. For co-immunoprecipitation analysis, 600 μg each of nuclear extracts were incubated with 10 μg of rabbit anti-Setd1a antibody (Bethyl) or 10 μg of normal rabbit IgG (Jackson ImmunoResearch Inc.) at 4°C, overnight. The immuno-complex was adsorbed onto protein A-agarose (Sigma) beads by incubating one hour at 4 °C. For immunoblot analysis we used rabbit anti-Mef2a/c antibody (1:1000, Abcam) or anti-Setd1a antibody (1:500) as a primary antibody and mouse monoclonal anti-rabbit IgG light chain (HRP) (1:5000 and 1:8000, clone SB62a, ab99697, Abcam) as a secondary antibody.

#### Immunocytochemistry

Primary neurons were fixed in 4% paraformaldehyde (PFA) in phosphate-buffered saline (PBS) (pH 7.4) for 10 min at room temperature (RT), washed 3 times for 5 min in 1×PBS, permeabilized in 0.1% Triton X-100 in PBS for 5 min and washed again in PBS (3 × 5 min). Coverslips were blocked for 30 min in 3% bovine serum albumin supplemented with normalized serum and incubated for 1 hr in primary antibodies diluted into blocking buffer and washed in PBS (3 × 5 min). Prior to application of secondary antibodies, cells were blocked again in blocking buffer supplemented with 0.5M NaCl for 30 min at RT. Secondary antibodies were applied in blocking/0.5M NaCl for 30 min at RT. Following another round of washing, the coverslips were mounted onto SuperfrostPlus glass slides using Prolong Gold (Invitrogen).

#### Immunohistochemistry

Experiments utilized brains from P8.5 or 6-weeks old males, perfused with PBS and 4% PFA, and post-fixed in PFA overnight. A vibratome was used to cut 60 µm coronal sections. Free-floating sections were blocked and permeabilized overnight at 4°C in PBTX (PBS/Triton X-100) containing 5% normal goat serum (PBTX / 5% NGS). Primary antibodies were diluted in PBTX / 5% NGS and incubated with sections for 24 hrs. at 4°C. Sections were washed overnight at 4°C in 1× PBS. Sections were incubated with secondary antibody (1:500, diluted in PBTX / 5% NGS) for 4 hrs. at RT. Following another round of washing in 1× PBS, sections were incubated with TO-PRO3 (1:2500 in PBS) for 30 min, washed in 1× PBS (4 × 15 min), and mounted on slides using Prolong Gold (Invitrogen).

#### Primary neuronal cultures and transfection

Primary cultures were prepared from cortices dissected bilaterally from E17 embryos. Cortices were individually dissociated using fire-polished Pasteur pipettes, counted, and plated in DMEM supplemented with 0.5% penicillin/streptomycin and 10% FBS. Neurons for immunocytochemical analyses were plated on poly D-lysine coated 12 mm coverslips (2 × 10^4^ cells/coverslip). Following 1.5 hr incubation, the modified DMEM was replaced with neurobasal media supplemented with B27 and Glutamax (Invitrogen) in which neurons were cultured for 5 days. Post-hoc genotyping was performed on DNA extracted from tissue taken from each embryo. Transient transfections were carried out with Lipofectamine 2000 (Invitrogen) at DIV4 using the protocol suggested for neuronal transfection. The original media used to culture cells prior to transfection were replaced in each well/dish following 2 hr of incubation at 37°C with transfection reagents.

*Administration of Kdm inhibitors into primary neuronal cultures.* KDM inhibitors were dissolved at 10 mM in 100% DMSO. The compounds were diluted to 0.1 nM - to 30 μM in fresh neurobasal media supplemented with B27 and Glutamax. Primary neuronal cultures were placed in neurobasal media with compounds at DIV3 and cultured for 48 hours. Transient transfections of pCAG-GFP were carried out with Lipofectamine 2000 at DIV4. Media with compounds used to culture cells prior to transfection were replaced in each well/dish following 2 hr of incubation at 37°C with transfection reagents.

#### Electroporation *in utero*

Pregnant mothers with E15.5 embryos were anaesthetized, and a 1 inch incision was made and one uterine horn removed. A total of 0.1–0.5 ml DNA (1–2 mg/ml) was injected into one ventricle. Four to five 40–45 V pulses were applied, each pulse lasting 50 ms, with a 1 s pause between each pulse using an ECM-830 square wave electroporator (BTX) as previously described (Mukai et al., 2015; Saito, 2006).

#### Virus injection

rAAV2/hSynapsin/EGFP was obtained from Addgene. Viral titer was 3 × 10^12^ particles per ml. Mice (8-12 week old) were deeply anesthetized under isoflurane and placed in a stereotaxic frame. For analysis of contralateral axon terminal branching *in vivo*, Bregma and lambda were used for leveling and zeroing, and bilateral cranial windows were excised over S1 cortex, centered at 0.97 mm posterior, 2.9 mm lateral. Virus was administered to layer 2/3 by an injection pump at a rate of 100 nL/min via an elongated 10-40 mm-diameter glass micropipette. 200 nl were infused at single injection site. For morphological analysis of spines in L2/3 neurons, Bregma and lambda were used for leveling and zeroing, and bilateral cranial windows were drilled over prefrontal cortex, centered at 1.93 mm anterior, 0.25 mm lateral. Virus was administered to L2/3 by an injection pump at a rate of 100 nL/min via an elongated 5-10 μm-diameter glass micropipette. 500 nl were infused at single injection site. Acrylic dental cement was used to cover the craniotomy after infusion.

#### Electrophysiology in medial PFC slice preparations

Field recordings were performed on 8- to 10-week-old male *Setd1a^+/-^* mutant mice and their WT littermates. Isoflurane was used to anesthetize mice that were then decapitated. After a skull incision, the brain was removed and placed in ice-cold dissecting solution, which contained 195 mM sucrose,10 mM NaCl, 2.5 mM KCl, 1 mM NaH_2_PO_4_, 25 mM NaHCO_3_, 10 mM glucose, 4 mM MgSO_4_, and 0.5 mM CaCl_2_ and was bubbled with 95% O_2_/5% CO_2_ mixture. Coronal brain slices (300µm) containing the prelimbic and infralimbic prefrontal cortex were cut using a vibratome (VT1200S; Leica, Nussloch, Germany). The freshly cut mPFC slices were immediately transferred to an interface chamber and allowed to recover for at least 2 h at 34–36°C. During all recordings, the slices were continuously perfused with artificial cerebrospinal fluid (aCSF) (bubbled with 5% CO_2_/95% O_2_) that had the following composition: 124 mM NaCl, 2.5 mM KCl, 1 mM NaH_2_PO_4_, 25 mM NaHCO_3_, 10 mM glucose, 1 mM MgSO_4_, and 2 mM CaCl_2_. The aCSF was maintained at 34–36°C and fed by gravity at a rate of 2–3 ml/min. Field EPSPs (fEPSPs) were recorded via a glass microelectrode (3–5 MΩ) filled with aCSF and placed in L5 of the mPFC (600–700 μm from midline). The stimulation site was always aligned ∼200 μm away from the recording site along the axis perpendicular to the pial surface. Basic synaptic transmission was characterized at 0.033 Hz, with stimulation intensities of 1–24 V (pulse duration, 0.1 ms). Subsequent experiments were performed at the stimulus intensity that generated a fEPSP one-third of the maximum fEPSP obtained at 24 V. Short-term synaptic facilitation was induced using a paired-pulse protocol with ISIs of 20, 50, 100, 200, 400, and 800 ms. To assess STD, fEPSPs were evoked by using a 40-pulse train at 5, 20 and 50 Hz (pulse duration, 0.1 ms). LTP was induced by one 800-ms (40 pulses), 50-Hz train after a stable 10 min baseline and monitored during 15 min. Then, 15 min after the first tetanus, four additional 50 Hz trains (separated by 10 s) were applied. The fEPSPs were then monitored for 40 min. Fiber volley was quantified by measuring the amplitude of the first peak negativity of the field responses, and the fEPSPs were quantified by measuring the initial slope of the second peak negativity of the responses. Electrophysiological signals were acquired using an extracellular amplifier (Cygnus Technologies) and pClamp 10 (Molecular Devices) Whole cell recordings were performed on 8- to 10-week-old male *Setd1a^+/-^* mutant mice and their WT littermates. Following brain slicing mPFC slices were immediately transferred to a recovery chamber and incubated at room temperature in recording solution (ACSF) for a minimum of 1h before recording. At the time of recording, slices were transferred to a submerged recording chamber and continuously perfused with ACSF with the temperature maintained between 30°C-34°C. Whole-cell patch clamp recordings were made using borosilicate glass pipettes (initial resistance 3.0–5.5 MΩ). Series resistance (5–20 MΩ) was monitored throughout each experiment and cells exhibiting a >20% change in series resistance were discarded. The internal solution used contained (in mM): KMeSO_4_ 145, HEPES 10, NaCl 10, CaCl_2_ 1, MgCl_2_ 1, EGTA 10, MgATP 5, Na2GTP 0.5, pH 7.2 with KOH, adjusted to 290 mOsm with sucrose. Layer 2/3 pyramidal neurons in mPFC prelimbic and infralimbic regions were identified by morphology and only neurons exhibiting spike frequency accommodation within action potential trains were included in analysis (Madison and Nicoll, 1984). All data were acquired with a Multiclamp 700B amplifier (Molecular Device) in conjunction with pClamp 10 software at 10kHz and low-pass filtered (10 kHz Bessel).

For recordings assessing neuronal excitability, action potential firing was assessed in current-clamp mode in response to incremental (20 pA steps), depolarizing current injections of 500 ms duration. Bridge balance of series resistance was employed and recordings with series resistance >20 MΩ were rejected. The resting membrane potential was determined within the first 5 s of achieving whole-cell access. For current-step experiments, the resting membrane potential of all cells was adjusted to approximately -70 mV by injection of a small standing current. Input resistance was calculated in current-clamp mode at -70 mV from the quasi steady-state (after 450-500 ms) voltage response to a 40 pA hyperpolarizing current-step. The membrane time constant (τ_memb_.) was calculated by fitting a single exponential to the voltage response to hyperpolarizing current steps (for 20-60 pA steps, using the voltage response during the first 200 ms) employing the built-in Chebyshev routine in Clampfit. For action potential afterhyperpolarization (AHP) analysis, the 1^st^ current step sweep producing at least two APs was used. As the interval between the 1^st^ and 2^nd^ APs was often very brief and complex (with the frequent presence of a “contaminating” after-depolarization waveform), the AHP after the 2^nd^ AP was used for analysis. The AHP was quantified as the difference between the AP threshold voltage and the minimal trough value of the AHP. The current-step trace used for analysis was overlaid with its own dV/dt plot to enable more confident assignment of AP threshold voltages.

For recordings assessing voltage-gated channel activation in response to voltage steps, cells were held at -70mV, 10mV steps were applied, and series resistance-related errors were partially corrected by using 70% prediction and 70% series resistance compensation. Voltage-activated currents are reported as the quasi steady-state currents at the end of a 100 ms voltage step.

#### Behavior analysis

##### Open field

Mouse activity was monitored in an illuminated chamber equipped with infrared sensors to automatically record horizontal and vertical activity (Colbourn Instruments). Each mouse was placed in the center of the chamber and its activity was recorded and collected in 1-min bins for 1 h during day 1 and for a 30-min re-exposure 24 h later (day 2). *T-maze:* 6 weeks old animals were trained on a non-match to sample T-maze task. The behavior protocol began with 2 days of habituation during which animals freely explored the maze for 10 minutes, followed by 2 days of 28 shaping when animals were required to alternate between goal arms of the maze to receive food rewards. After shaping was completed, training and testing sessions were conducted. Each trial of the task consisted of a sample and choice phase. In the sample phase, a mouse ran down the center arm of the maze and was directed into one of the goal arms. The mouse returned to the start box where it remained for a delay of 10 seconds. In the choice phase, the mouse was required to select the arm opposite to that visited in the sample phase to receive a reward. Animals were given daily training of 10 trials until they reached criterion performance, defined as performance of at least 70% correct per day for 3 consecutive days. After criterion was reached, animals completed daily testing sessions composed of 20-25 trials for delays of 10 s, 30 s, and 60 s. In the case of rescue experiments following treatment with tamoxifen, TCP, or ORY-1001, animals completed daily testing sessions composed of 12 trials for 10 s, 20 s, and 40 s delays.

##### Social memory-Direct interaction with juveniles

All mice were housed two to five in each cage and given *ad libitum* access to food and water. They were kept on a 12-h (6:00 to 18:00) light–dark cycle with the room temperature regulated between 21-23 °C. Behavioral tests were performed during the light cycle in a testing room adjunct to the mouse housing room, which minimizes any transportation stress. The social memory/direct interaction test with juveniles was adapted from previously described (Kogan et al., 2000). Immediately prior to the experimental sessions, 10-12-week-old *Setd1a^+/-^* and WT littermates were transferred to the testing room and placed into individual cages, identical to the ones used for housing, where they were allowed to habituate to the new environment for 15 min. Male juvenile mice (C57BL/6J, 4-5-week-old) were also placed in the testing room in their home cages and allowed to habituate a similar amount of time. Testing began when a novel juvenile mouse was introduced to a cage with one of the adult experimental mice. Activity was monitored for 5 min (trial 1) and scored online by a trained observer blind to the genotype of the test mice for social behavior (anogenital and nose-to-nose sniffing, close following and allogrooming) initiated by the. experimental subject, as described in Hitti et. al. (Hitti and Siegelbaum, 2014). After an inter-trial interval of 1 h, the experimental mice were introduced with either the previously encountered mouse or a novel mouse again for 5 min (Trial 2). The time spent in social interaction during trial 1 was subtracted from the social interaction time during trial 2 to obtain the difference score.

##### Fear Conditioning

Experimentally naive mice were used for fear conditioning in plastic, sound-attenuating cabinets containing a house light. On the 1st day, mice were placed in chambers and received 2 pairings of a tone (30 s, 82 db) and a co-terminating shock (2 s, 0.7 mA). On the 2nd day, initially, mice were placed in the chambers for 6 min, in the absence of tone and shock, and freezing was scored for the whole session, corresponding to hippocampal-dependent contextual fear memory. 2 h later, the context and handling of the mice was changed to assess conditioned fear of the tone alone (cue fear memory). Freezing was assessed before and during the tone presentation to distinguish fear generalization and specific cue fear memory for cue.

##### Novel Object Recognition

Mice were given a single 10 min trial of habituation to the room, the testing arena and to the procedure over three consecutive days to reduce stress and prevent a neophobic response and consequently to promote the exploratory activity of mice toward the objects (days 1-3). On the two following days mice were given a 5 min trial of exposure to two identical objects (days 4, 5). A memory test trial was performed 1 h later where one of the familiar objects was replaced with a novel object. 24 h later mice were placed back in the arena and were given another 5 min memory test in which one of the identical objects they explored during the training trial was replaced with a novel one. The location of the novel object was randomized between trials and genotypes for both the 24 h and the 1 h novel object recognition test. A minimal exploration time for both objects during the test phase (∼20 s) was used to ensure a similar exploration time of the two objects and between animals, independently of their individual exploratory activity. All trials were video recorded and analyzed blindly by two investigators. Results were expressed as a discrimination index, calculated as the amount of time each mouse spends exploring the novel object divided by the total exploration time for both objects.

#### Tamoxifen preparation and feeding

We used a modified version of a tamoxifen feeding protocol previously described in Mei, et al (Mei et al., 2016). Tamoxifen (Sigma, T5648) was dissolved in corn oil (Sigma, C8267) at 20 mg/ml by vortexing. Freshly prepared tamoxifen was protected from light by aluminum foil. Animal feeding needles (Harvard Apparatus, 52-4025) were used for oral gavage. To avoid toxicity of tamoxifen, the following dosages were used for adult animals (8 weeks): mice at 17–21 g body weight were fed 5 mg/day; mice at 22–25 g body weight were fed 6 mg/day; mice at 26–29 g body weight were fed 7mg/day; mice at 30–35 g body weight were fed 8mg/day. Adult animals were fed for 5 consecutive days followed by 2 days of rest. Animals were then fed for 5 more consecutive days followed by 3 weeks of rest. Corn oil was used as a control. Mice fed with tamoxifen and with corn oil were housed separately to avoid contamination.

#### TCP preparation and injection

Tranylcypromine (TCP, Sigma, P8511) was dissolved in PBS at 0.75 mg/ml by vortexing. Mice at 17-20 g body weight were injected 2.7 mg/kg/day; mice at 20-28g body weight were injected 3.0 mg/kg/day. Adult animals were injected for 14 consecutive days followed by a non-match to sample T-maze task. During the behavior task, mice were injected with the same dose at 6 pm daily.

#### ORY-1001 preparation and injection

ORY-1001 (Cayman Chemical, 19136) was dissolved in PBS at 1mg/ml by vortexing. Mice at 17-20 g body weight were injected 9.0 μg/kg/day; mice at 20-28g body weight were injected 10.0 μg/kg/day. Adult animals were injected for 14 consecutive days followed by a non-match to sample T-maze task. During the behavior task, mice were injected with the same dose at 6 pm daily.

#### RNA sequencing

Total RNA was extracted from prefrontal cortices of 6-week-old mice using RNeasy Mini Kit (Qiagen). We analyzed each four male *Setd1a^+/-^* mice and their wild-type (WT) male littermates. Quality of RNA samples was assessed by Bioanalyzer (Agilent). Mean (± SD) RNA integrity number (RIN) was 9.3 (± 0.5) in the mutant mice and 9.7 (± 0.2) in WT. An RNA-seq library was prepared from ∼400 ng of total RNA by using TruSeq RNA prep kit (Illumina). Messenger RNAs were enriched by poly-A selection. Libraries were sequenced on HiSeq2500/HiSeq4000 (Illumina) with single-end 100bp reads at Columbia Genome Center.

#### ChIP-Seq analysis

Whole prefrontal cortex was dissected from 6-week old wild-type (WT) mice. Dissected tissue was thoroughly minced on ice using razor blades, then fixed with EGS for 30 min and/or for 5 min at room temperature with methanol-free formaldehyde (Pierce, Thermo Fisher Scientific) diluted to 1% in PBS. Fixation was quenched by adding 1/10th volume of 1.25 M glycine. Fixed tissue was collected by centrifugation at 200 g for 5 min at 4°C, then washed twice with cold PBS. Washed tissue was flash-frozen with liquid nitrogen then cryofractured using a Covaris CryoPrep Impactor on power setting 6. Nuclei and chromatin immunoprecipitation were performed as in Markenscoff-Papadimitriou *et al*.(Markenscoff-Papadimitriou et al., 2014): briefly, the cryofractured tissue was lysed in ChIP Lysis Buffer (50 mM Tris-HCl pH 7.5, 150 nM NaCl, 0.5% NP-40, 0.25% Sodium Deoxychoalate, 0.1% SDS) in rotation for 40 min at 4°C. Nuclei were centrifuged (1700x g, 5 min, 4°C), and the pellet was resuspended in shearing buffer (10 mM Tris-HCl pH 7.5, 1 mM EDTA pH 8, 0.25% SDS), then sheared on a Covaris S2 sonicator (12 or 16 min, 2% Duty Cycle, Intensity 3, 200 cycles per burst, frequency sweeping). Sheared chromatin was centrifuged (10,000 g for 10 min at 4°C) to remove insoluble material. Sheared chromatin was diluted 5-fold with ChIP Dilution Buffer (CDB: 16.7 mM Tris-HCl pH 8.1, 167 mM NaCl, 1.2 mM EDTA, 1.1% Triton X-100, 0.01% SDS) and pre-cleared for two hours at 4°C with protein G dynabeads (Thermo Fisher Scientific). Each ChIP was set up with the appropriate antibody [10 μg of anti-Setd1a (hSET1, Abcam, ab70378), 1 μg of anti-H3K27ac, -H3K4me2 and -H3K4me3, 3 μg of anti-H3K4me1, 5 μg of anti-Mef2] and 10-20 μg of cleared chromatin, incubating then overnight at 4°C. Protein G dynabeads were blocked overnight with 2 mg/ml yeast tRNA (Thermo Fisher Scientific), then added to antibody bound chromatin and rotated for 3 hr at 4°C. Bead-bound chromatin was washed 5 times with LiCl Wash Buffer (100 mM Tris-HCl pH 7.5, 500 mM LiCl, 1% NP-40, 1% Sodium Deoxycholate) and once with TE (pH7.5). DNA was eluted from beads by incubating at 65°C for 30 min with 25 μL ChIP Elution Buffer (1% SDS, 0.1 M Sodium Bicarbonate). This elution was repeated and the combined elution fractions were incubated overnight at 65°C with proteinase K (Thermo Fisher Scientific) and RNaseA (Thermo Fisher Scientific). ChIP DNA was purified 1.8X of AMPure XP beads and eluted in 30 μl of nuclease free water. Libraries were prepared with Ovation Ultralow System V2 1-16 (NuGEN technologies) according to manufacturers’ recommendation. All samples were multiplexed (in batches of 6) and sequenced using Illumina HiSeq as 50 bp or 75 bp single end. Two biological replicates for each ChIP-Seq for each transcription factor or histone mark were sequenced. As control, 1% chromatin input was also sequenced.

#### Dissociation of PFC to single cells and scRNA-Seq analysis

Cells were dissociated using the papain dissociation kit (Worthington) following the manufacturers’ recommendation. In brief, the PFC of 2 WT and 2 *Setd1a^+/-^* mice was dissected and transferred to ice-cold PBS. The PFC was cut in small pieces using a razor blade and dissociated with a papain dissociation system (Worthington Biochemical). The tissue was incubated for 1h at 37°C on a rocking platform with papain-EBSS (1 PFC/mL). The tissue was triturated 20 times and cells were pelleted (400 g, 5 min, RT). Remaining papain was inhibited by resuspending the cell pellet with Ovomucoid protease inhibitor solution diluted 1:10 in EBSS. An extra 5 ml of Ovomucoid protease inhibitor solution was slowly layered at the bottom of the cell suspension to create a discontinuous gradient and cells were pelleted (100 g, 6 min, RT). To enrich for living cells, we removed dead cells from single cell suspension using the MACS® Dead Cell Removal Kit (Miltenyi Biotec) with the MS Columns following manufacturer’s recommendation. In brief, 200 μl per PFC of Dead Cell Removal MicroBeads were added the pelleted cells. Cell were resuspended and incubated for 15 min at RT. The MS columns were washed once with 0.5 ml of Binding buffer. After incubation is complete, the cell suspension was diluted with 500 μl 1X Binding Buffer and the suspension was applied to the MS columns. The flow-through was collected. To increase the number of live cells recovered, the column was washed twice with 1 ml of Binding buffer at RT. Flow-through were combined and live cell were pelleted (400 g, 5 min, RT). Cells were then washed twice with PBS containing 0.04% BSA and the final pellet was resuspended in 300 μl of PBS containing 0.04% BSA and passed through a 40 uM cell strainer. Cell were counted and viability was evaluated by trypan blue staining.

The scRNA-Seq was performed using the 10x Genomic platform and the “Single Cell 3’ v2” by the Columbia Genome Center. The scRNA-Seq data were aligned using the 10x Genomics package Cell Ranger 2.1.1 on the mm10 mouse genome version. In particular, UMI reads from the different genotypes and biological replicates were counted using “cellranger count” and combined together using “cellranger aggr”. From the downstream analysis, we removed cells in which we detected less then 1000 genes expressed, using “Seurat” R-package (v. 3.0.1). This package was used for all downstream analysis. In details, counts were first normalized (using the normalization method: “LogNormalize”). Linear dimensionality reduction was then performed on the most variable genes after that the data were regress out for cell-cell variability (i.e. batch effect, cell alignment rate, number of detected molecules etc.). All significant principal components were calculated, selected (using 40 PCs) and used to defined cell clusters (with the function “FindClusters”). Clusters were then visualized through UMAP plots using the same number of PCs that were used to define clusters. The different cell types were finally defined by the known expression of genes that have been previously describe to mark specific subpopulation of neuronal and non-neuronal cells (Figures S6B, S6C, and S6D). Genes with average expression < 0.1 normalized UMI counts in the whole dataset were removed from all analyses performed.

#### Sorting of PFC pyramidal neurons and parvalbumin expressing interneurons to identify hypersensitive genomic regions

Emx-Cre mice (Jackson lab) and Parv-Cre mice (Jackson lab) were crossed to CRE inducible tdTomato mice (Jackson lab) so that pyramidal neurons or parvalbumin expression interneurons expressed the fluorescent protein tdTomato. The PFC of these mice was dissociated as described previously. After dissociation cells were resuspended in sort media supplemented with 100 U/mL DNase I (Worthington Biochemical), 4 mM MgCl_2_, and 500 ng/mL DAPI (Invitrogen). These cells were passed through a 40 uM cell strainer, and then FAC sorted. Live cells were selected by gating out DAPI positive cells and positive cells were gated by tdTomato fluorescence. Fifty thousand cell were used to prepare ATAC-seq libraries as described in references (Buenrostro et al., 2015; Monahan et al., 2017). In brief, cells were pelleted (500 g, 5 min, 4°C) and resuspended in ice cold lysis buffer (10 mM Tris-HCl, pH 7.4, 10 mM NaCl, 3 mM MgCl_2_, 0.1% IGEPAL CA-630). Nuclei were pelleted (1000 g, 10 min, 4°C) and resuspended in transposition reaction mix prepared from Illumina Nextera reagents (for 50 μL: 22.5 μL water, 25 μL 2xTD buffer, 2.5 μL Tn5 Transposase). Transposed DNA was column purified using a Qiagen MinElute PCR cleanup kit (Qiagen) and amplified using barcoded primers and NEBNext High Fidelity 2x PCR Master Mix (NEB). Amplified libraries were purified using Ampure XP beads (Beckman Coulter). ATAC-Seq data were aligned and processed as previously described for ChIP-Seq data.

#### Luciferase assay

HEK293T cells were transfected in a 96well plate using Lipofectamine 3000 (ThermoFisher Scentific) following the manufacturer’s instruction. We transfected 5ng of pRL Renilla vector, 50 ng of 3X Mef2-Luciferase vector (Addgene #32967), 75 ng of Mef2c CDS (Addgene #32515) and/or Setd1a CDS (Origene #MR215352), Lsd1 CDS (Origene #MR210741), pcDNA3.1 vector. Luciferase assay was performed using the kit Dual-Glo® Luciferase Assay System (Promega, #E2920) following the manufacturer instructions. Measurements were performed using the plate reader SpectraMax M3 (Molecular Devices).

### QUANTIFICATION AND STATISTICAL ANALYSIS

#### Quantification of axon branching, dendritic spines, dendritic complexity and neuronal numbers

Quantification, image and statistical analyses cytoarchitecture were performed with LSM Image browser tool, Fiji (Schindelin et al., 2012) and Excel software, blind to the genotype. Quantifications throughout the study are depicted as mean values with error bars indicating the Standard Error of the Mean (s.e.m). The number of independent repeats (N), the statistical test used for comparison and the statistical significance (P values) are specified for each figure panel in the representative figure legend. P values in this manuscript present the following star code: P>0.05 (non-significant), *P < 0.05, **P < 0.01, ***P < 0.001.

##### Quantification of axonal morphology in vivo

To analyze the axonal morphology quantitatively, we performed the following procedures for each dataset obtained from the 60-μm coronal sections. For the contralateral axon branch number at P8.5, images were taken with setting as z-series stacks (at 0.5 µm depth intervals) using 40× Zeiss objectives with constant parameters for brightness and contrast. EGFP expressed axons at contralateral layer 1-4 were analyzed by projecting the *z* series and manually tracing the axon using the overlay tools of the LSM Image Browser program (Zeiss). Protrusions more than 6 μm long were categorized as axon branches. The axon terminal architecture of each neuron was also characterized by designating the order of a branch (primary, secondary and so on). Once the elaborations of the axon branch tree were outlined, branch numbers were counted manually and analyzed. We compared at least 20 axons from four brains of each genotype in three independent experiments using the Student’s *t*-test.

##### Quantification of cortical axonal morphology in vivo in adult mice

To analyze and visualize axon branching in adult brain, we performed the following procedures for each dataset obtained from 300-μm coronal sections. Images acquired for the EGFP fluorescence analysis were taken with sequential acquisition setting at 2,048 × 2,048 pixel resolution as z-stack (taken at intervals of 1.5 μm) from each sections of frontal cortex using Nikon eclipse Ti2 + W1-Yokogawa spinning disk confocal microscope. Z-stack images for whole contralateral axon column in the somatosensory cortex (S1) were projected by z-project mode with Sum Slices projection type of ImageJ Fiji software. The immunofluorescence in each layer was measured by ImageJ Fiji software and normalized by EGFP expressed cell number in ipsilateral layer 2/3 as described below. All settings of threshold were maintained constant across genotypes. To quantify the number of cells expressing EGFP at the injection site of ipsilateral S1 cortex, images were taken with sequential acquisition setting at 2,048 × 2,048 pixel resolution as z-stack (taken at intervals of 2.0 mm) from each sections of frontal cortex using Nikon eclipse Ti2 + W1-Yokogawa spinning disk confocal microscope. The number of cells expressed EGFP were counted using 3D Objects Counter of ImageJ Fiji software.

##### Quantification of axonal morphology in vitro

For *in vitro* analysis, a plasmid, pCAG-GFP, expressing EGFP driven by the β-actin promoter was transfected into primary neuron cultures to fill the cells and facilitate visualization of all processes. Confocal images of neurons from primary culture were obtained blind to genotype with the LSM510 using a Zeiss 20× objective with sequential acquisition setting at 2048 × 2048 pixels resolution. Images were taken with setting as z-series stacks (at 1.0 µm depth intervals) with constant parameters for brightness, contrast, and pinhole size. Neuronal processes stained for the axonal marker Tau-1 were traced using the overlay tools of the LSM Image browser program (Zeiss). Protrusions more than 8 μm long were categorized as axon branches. The axon architecture of each neuron was also characterized by designating the order of a branch (primary, secondary and so on). Once the elaborations of the axon branch tree were outlined, branch points were counted manually, and the length of primary axon (emanating directly from the soma) and its branches were analyzed. To evaluate the axon length and branching number, all EGFP-expressed cell, which were polarized with single axon (approximately 50 - 80 neurons/coverslip), were analyzed from at least 5 coverslips derived from 3 embryos for each genotype. using the Student’s *t*-test.

##### Quantification of dendritic complexity

To analyze the dendritic complexity of the basal dendrites of L5 neurons in the PrL area of Thy1-GFP/M mice at 6-weeks, images were taken with setting as z-series stacks (at 0.5 µm depth intervals) using 40x Zeiss objectives with constant parameters for brightness and contrast. The complexity of the basal dendrites was analyzed by projecting the z-series to maximum intensity and manually tracing the dendritic tree. Sholl analysis was conducted using concentric radii of 20 mm extending from the center of the soma. Intersections, branches and terminals were counted by distance from the soma and differences evaluated by repeated measures ANOVA.

##### Quantification of dendritic spines

To analyze the dendritic spine number, images were taken with setting as z-series stacks (at 0.3 µm depth intervals) using 60x Zeiss objectives with constant parameters for brightness and contrast. Z-series stacks were subjected to three-dimensional reconstruction using Zeiss LSM 510 Meta. Spine counts were performed on the second dendritic branch of intact neurons beginning at 25–40μm away from the soma and extending for 30–60μm from the origin. Dendritic protrusions were classified as spines (mushroom, long, stubby or thin) or filopodia, based on the parameters previously outlined (Mukai et al., 2008). Differences in the number of spines were evaluated using the Student’s *t*-test. Size determination of mushroom spines was carried out using Zeiss LSM image browser overlay tools. The longest straight line in the spine head was counted as the head diameter. The length of the entire spine (including head and stalk) was measured using a bent-line tool and analyzed by Kolmogorov-Smirnov test.

##### Quantification of neuronal numbers in tissue sections

To quantitate the NeuN and PV positive cell number at 6-weeks, images were taken with setting as a single scan using 20× Zeiss objectives with constant parameters for brightness and contrast. To analyze the VIP-, SST-, and Reelin-positive cell number at 6-weeks, images were taken with setting as z-series (at 3 µm depth intervals) using 20x Zeiss objectives with constant parameters for brightness and contrast.

#### Electrophysiological analysis

Statistical analyses were performed using SigmaPlot 9.0, Microsoft Excel and GraphPad Prism 4. A two-way repeated-measures ANOVA was used to test differences between genotypes. For simple comparisons of single metrics, unpaired t-tests were employed. For t-tests, if there was a directional hypothesis, 1-tailed t-tests were performed. For AP responses across current steps, 2-way repeated measures ANOVA statistics were employed. In the corresponding figures, data are presented as means ± s.e.m. N indicates number of animals and n indicates number of slices or cells from which recordings were made. Asterisks indicate significant differences. All recordings and the majority of data analyses were done blind to the genotype.

#### Behavioral analysis

Statistical analyses were performed using SPSS 20.0 and GraphPad Prism 6. Statistical analyses for behavioral tests were performed using the Student’s two-tailed *t-*test (T-maze, open field, novel object recognition memory, and fear conditioning) or two-way ANOVA and post hoc pairwise comparisons with Bonferroni correction (social memory). Quantifications throughout the study are represented as the mean value. Error bars indicate the Standard Error of the Mean (s.e.m).

#### Data processing and analysis of DE genes

We used RTA (Illumina) for base calling and bcl2fastq (Illumina, version 2.17) for converting BCL to fastq format, coupled with adaptor trimming. We mapped the reads to the reference mouse genome (UCSC/mm10) using STAR ((Dobin et al., 2013) (version 2.5.4b). Read counts and normalization of the read count data and analysis of differentially expressed genes (DE genes) were performed by using DESeq2 (Love et al., 2014) (version 1.10.2) with default parameters. By performing principle component analysis (PCA) with our data of eight PFC samples (each five *Setd1a+/-* and WT) and read count data from olfactory bulb (OB) and cerebellum, we confirmed that our data of PFC form a single cluster distant from clusters of OB and cerebellum (data not shown).

#### ChIP-Seq data processing, peak calling and motif discovery

Reads were aligned to the *Mus musculus* genome version mm10 using bowtie2 v2.2.6 (Langmead and Salzberg, 2012). Poor quality and duplicated reads were removed using respectively samtools v1.3.1(Li et al., 2009) and Picard (http://broadinstitute.github.io/picard). To measure the reproducibility between biological replicates and to avoid arbitrary thresholding biases for both Setd1a and Mef2, we perform IDR following the ENCODE framework (Landt et al., 2012) using MACS v2.1.1 as peak caller (DOI: 10.1186/gb-2008-9-9-r137) and IDR 2.0.2 (Li et al., 2011). We defined good biological replicates when the ratio N1/N2 and Np/Nt was below 2. For all subsequent analysis, biological replicates were merged into single alignment files for each antibody using samtools v1.3.1 (Li et al., 2009) and difference in sequencing depth between the libraries was corrected by normalization for library size using HOMER v4.8.2 (Heinz et al., 2010). Setd1a bound promoter elements were defined as such when they were located within 1 Kb of annotated TSS or they overlapped with H3K4me3; Setd1a bound intergenic peaks were defined as such when they were located in unannotated regions of the genome at least 1 kb away from any annotated TSS; Setd1a bound intronic peaks were defined as such when they overlapped with annotated intron of the *Mus musculus* (version mm10) of the genome. *De novo* motif discovery analysis was performed on feature-separated *Mus musculus*, version mm10, of Setd1a peaks using HOMER v4.8.2 (Heinz et al., 2010) using automatically generated background (sequences randomly selected from the genome, matched for GC% content).

#### Setd1a binding in promoter, intergenic and intronic regulatory regions

To identify the properties of Setd1a bound regions in the PFC, we compared the coverage of PFC ChIP-Seq for Mef2 and histone marks that we sequenced (see above). Heatmaps were generated calculating the normalized coverage per each ChIP-Seq using deeptools v2.5.0.1 (Ramirez et al., 2016), 2 kilobases around the center of Setd1a peaks into 50 bp bins. Density plots were generated calculating the coverage of the ChIP-Seq reads 2 kilobases around the center Setd1a peaks into 50 bps bins with deeptools v2.5.0.1 (Ramirez et al., 2016) and plotted using the R-package “ggplot2” and R v3.6 (http://www.gbif.org/resource/81287).

#### Brain region-specific Setd1a binding

To investigate the Setd1a chromatin binding pattern between different tissues, we compared our PFC epigenetic H3K27ac landscape with different brain tissues using the ENCODE publicly available H3K27ac ChIP-Seq data from the olfactory bulb and the cerebellum (Shen et al., 2012). To make the data analysis comparable between datasets, we reprocessed the ENCODE data as described above. Heatmaps were generated as described above.

#### Conservation of Setd1a peaks in the Euarchontoglire clade

Phastcons were obtained using the 59 vertebrate genomes (Phastcons 60ways) (Siepel et al., 2005). We selected data from the Euarchontoglire clade (including 21 vertebrate genomes) and the sources were downloaded from UCSC Genome browser via http://hgdownload.cse.ucsc.edu/goldenpath/mm10/phastCons60way/. Setd1a peaks were first normalized by size and heatmaps were generated as described above. Clusters were defined by hierarchical clustering of the data. For cluster 2 and 3 (Figure S3E), we split peaks in half: conserved and non-conserved half. Mef2 motif distribution was calculated using HOMER v4.8.2 (Heinz et al., 2010) with the function “annotatedPeaks” and plotted using the R package “ggplot2”. De novo motif analysis was performed with HOMER v4.8.2 (Heinz et al., 2010) automatic background.

#### Conservation of murine Setd1a peaks in the human genome

To evaluate the conservation of Setd1a peaks between mouse and human, we submit the FASTA sequence of 12499 PFC specific Setd1a peaks to the online blast platform https://blast.ncbi.nlm.nih.gov/Blast.cgi. We used the megablast algorithm (McGinnis and Madden, 2004) to look for very similar sequences (>95% identity) between the selected species. We found 3813 conserved-Setd1a-bound regions between mouse and human. We performed *de novo* motif analysis on the hg38 version of the human genome using HOMER v4.8.2 (Heinz et al., 2010) and automatic generated background.

#### scRNA-Seq analysis

The scRNA-Seq data were aligned using the 10x Genomics package Cell Ranger 2.1.1 on the mm10 mouse genome version. In particular, UMI reads from the different genotypes and biological replicates were counted using “cellranger count” and combined together using “cellranger aggr”. From the downstream analysis, we removed cells in which we detected less then 1000 genes expressed, using “Seurat” R-package (v. 3.0.1). This package was used for all downstream analysis. In details, counts were first normalized (using the normalization method: “LogNormalize”). Linear dimensionality reduction was then performed on the most variable genes after that the data were regress out for cell-cell variability (i.e. batch effect, cell alignment rate, number of detected molecules etc.). All significant principal components were calculated, selected (using 40 PCs) and used to defined cell clusters (with the function “FindClusters”). Clusters were then visualized through UMAP plots using the same number of PCs that were used to define clusters. The different cell types were finally defined by the known expression of genes that have been previously describe to mark specific subpopulation of neuronal and non-neuronal cells (Figures S6B, S6C, and S6D). Genes with average expression < 0.1 normalized UMI counts in the whole dataset were removed from all analyses performed.

#### ATAC-Seq analysis

ATAC-Seq data were aligned and processed as previously described for ChIP-Seq data.

#### Human ATAC-Seq data to evaluate the accessibility of Setd1a conserved regions

Downloaded ATAC-Seq data from postmortem human brain has been previously described (Fullard et al., 2018). The fastq file were realigned as described previosly (see ChIP-Seq analysis section of the methods) on the hg38 genome.

#### Luciferase assay analysis

Lucifarese values were normalized against Renilla values to account for differences in transfection efficiency. Three biological replicates (each having three technical replicates) on different batch of HEK293 cells were generated. Each condition was normalized on empty vector values. Values are display in Figure 5C and Figure 7D as mean ± standard error for three biological replicates. Student’s unpaired t-test was used to compare groups and test for significant differences.

#### Interval-based enrichment analysis of Set1a target genes

To enrich for functional elements in human, we identified conserved PFC-specific Setd1a peaks overlapping with ATAC-Seq from 5 different regions of the human brain (dorsal and ventral PFC, hippocampus, Medial dorsal Thalamus and putamen). By clustering these regions, we identified 1765 regions that are chromatin accessible in the human brain (Figure S4E). Finally, we used a permutation test implemented RegioneR (Gel et al., 2016) package in R to determine if there is any significant enrichment of these elements within genomic signals identified in previous GWAS studies of SCZ and other disorders.

#### Gene-based enrichment analysis of Set1a target genes among GWAS signals

For the gene-based enrichment analysis, all conserved, brain-region specific and chromatin accessible ChIP-Seq peaks were first annotated using the PAVIS annotation tool according to their relative positions to known genes in human genome. We then utilized the INRICH analysis tool (Lee et al., 2012), a permutation-based approach that controls for potential biases caused by SNP density, gene density and gene size to analyze the overlap between the human orthologue targets and previously reported GWAS signals to determine if there is any significant overlap. For the INRICH analysis, the reference gene list was downloaded from the INRICH website (https://atgu.mgh.harvard.edu/inrich/downloads.html). The reference SNP list of GWAS studies (SNPs on a DNA microarray) was compiled from the table browser (hg19.snpArrayAffy6 and hg19.snpArrayIlluminaHumanOmni1_Quad) of the UCSC genome browser, which covers at least 95% of intervals analyzed. All coordinates were converted to hg38 using the LiftOver tool at the UCSC browser (https://genome.ucsc.edu/cgi-bin/hgLiftOver). The default setting of 5000 permutation replicates was used for each trait tested. We prepared the various GWAS signals using the summary association data from the NHGRI (National Human Genome Research Institute) GWAS Catalog (Welter et al., 2014). Specifically, we first downloaded the list of SNPs associated with various human traits from the GWAS Catalog (http://www.ebi.ac.uk/gwas, the gwas_catalog_v1.0.1 file, accessed June 2016). For each phenotype, genome-wide significantly associated intervals were defined as follows using the threshold for linkage disequilibrium (LD) used in the SCZ GWAS by PGC (*r*^2^ > 0.6) (Schizophrenia Working Group of the Psychiatric Genomics, 2014). We extracted autosomal SNPs with genome-wide significant association (*P* < 5 × 10^-8^) and with MAF > 0.05 in the 1000 Genomes Project March 2012 EUR dataset [downloaded as a part of the LocusZoom package (Pruim et al., 2010)] from the full dataset of GWAS Catalog (we denote these as “index SNPs”). An associated genomic locus for each index SNP was defined as the genomic region containing any SNPs that are: 1) in LD with the index SNP at *r*^2^ > 0.6 and 2) within +/-10 Mb of the index SNP. SNPs in LD with the index SNP were identified by using PLINK (Purcell et al., 2007) with the 1000 Genomes Project March 2012 EUR dataset. As a control, we also performed the same analyses on GWAS signals of 15 non-CNS diseases/traits that have similar associated interval size (from 74 intervals to 174 intervals) that are comparable to the size of SCZ GWAS intervals. This procedure of generating independent GWAS signals is a pre-requisite for the INRICH analysis. INRICH analysis was conducted using version 1.1 with its default settings.

#### Gene-based enrichment analysis of Setd1a target genes among LoF DNM-hit genes

Gene-based enrichment analysis of the human target gene set of Setd1a (described above) among LoF DNMs was performed by using DNENRICH (Fromer et al., 2014), a statistical software package for calculating permutation-based significance of gene set enrichment among genes hit by de novo mutations accounting for potential confounding factors such as gene sizes and local trinucleotide contexts DNENRICH analyses were performed with one million permutations using the following input files: 1) gene name alias file: the default dataset included in the DNENRICH package, 2) gene size matrix file: the default dataset included in the DNENRICH package (for RefSeq genes), 3) gene set file: the set of 708 high-confidence Setd1a target genes (HGNC symbols available, not including *SETD1A*); all genes were equally weighted, 4) mutation list files: lists of genes hit by LoF DNMs in NDD (SCZ, ID/DD and ASD), congenital heart diseases (CHD) and control subjects in denovo-db (Turner et al., 2017) or Refs. (Xu et al., 2012; Xu et al., 2011; Yuen et al., 2017); all mutations were equally weighted, and 5) background gene file: the default gene list in the gene size matrix. The numbers of LoF DNMs in NDD, SCZ, ID/DD, ASD, CHD and controls were 1,874, 95, 986. 793, 173 and 202, respectively. For analyses restricting the input LoF DNMs to those in genes intolerant to LoF mutations, we used the two metrics, pLI scores (Lek et al., 2016) (threshold = 0.9) and LoF FDR (Petrovski et al., 2013) (threshold = 0.01). Genes for which pLI or LoF FDR was not available were excluded from the background gene list in the corresponding analyses considering pLI or LoF FDR.

#### GO enrichment and CSEA analysis of Setd1a target genes

Analysis of gene ontology (GO) terms enriched among Setd1a target genes was performed by using ToppGene with default parameters for four categories (Molecular Function, Biological Process and Cellular Component of GO terms, and Disease terms defined by DisGeNET). The enriched GO terms were visualized with REVIGO Gene Ontology treemap (Supek et al., 2011) (http://revigo.irb.hr/). To study the expression pattern of Setd1a targets, in the context human brain development, Setd1a target genes were imported into the CSEA tool (Dougherty et al., 2010) (http://genetics.wustl.edu/jdlab/csea-tool-2/). The output FDR *P* values of CSEA analysis were imported into the MultiExperiment Viewer (ver.4.9.0) as heatmap color code to generate the heatmap view. Genes that overlap with developmental stage and cortical signature genes were summarized from SEA analysis (pSI p< 0.05).

#### DATA AND CODE AVAILAVILITY

RNA-Seq (Figure 5 and Figure S6), scRNA-Seq (Figure 5 and S6) and ChIP-Seq (Figure 3, Figure S4 and Figure 7) raw and processed data are available at GEO repository (GSE123652). SCZ GWAS data is publicly available from http://www.med.unc.edu/pgc/results-and-downloads/. All other disease GWAS data is public available from http://www.ebi.ac.uk/gwas. LoF DNMs data is publicly available from http://denovo-db.gs.washington.edu/denovo-db/.

### ADDITIONAL RESOURCES

None

Supplemental Data (Excel files) Data Table 1

Setd1a promoter peaks (Related to Figure 3) Setd1a enhancer peaks (Related to Figure 3) Setd1a peaks all (Related to Figure 3)

Setd1a motif enrich enhancers (Related to Figure 3) Setd1a motif enrich promoters (Related to Figure 3)

hg38 megablast peaks accessible (Related to Figure 3, S3 and 4) motif megablast hg38 accessible (Related to Figure S3)

whole PFC RNA-Seq wt vs mut (Related to Figure S3)

#### Data Table 2

hg38 conserved accessible setd1a peaks (Related to Figure 4) PAVIS anno (Related to Figure 4)

intron (Related to Figure 4) allOthers (Related to Figure 4)

Not annotated (Related to Figure 4)

