## Supplementary Information for "Recapitulation and reversal of schizophrenia-related phenotypes in *Setd1a*-deficient mice"

#### Supplementary Figure 1

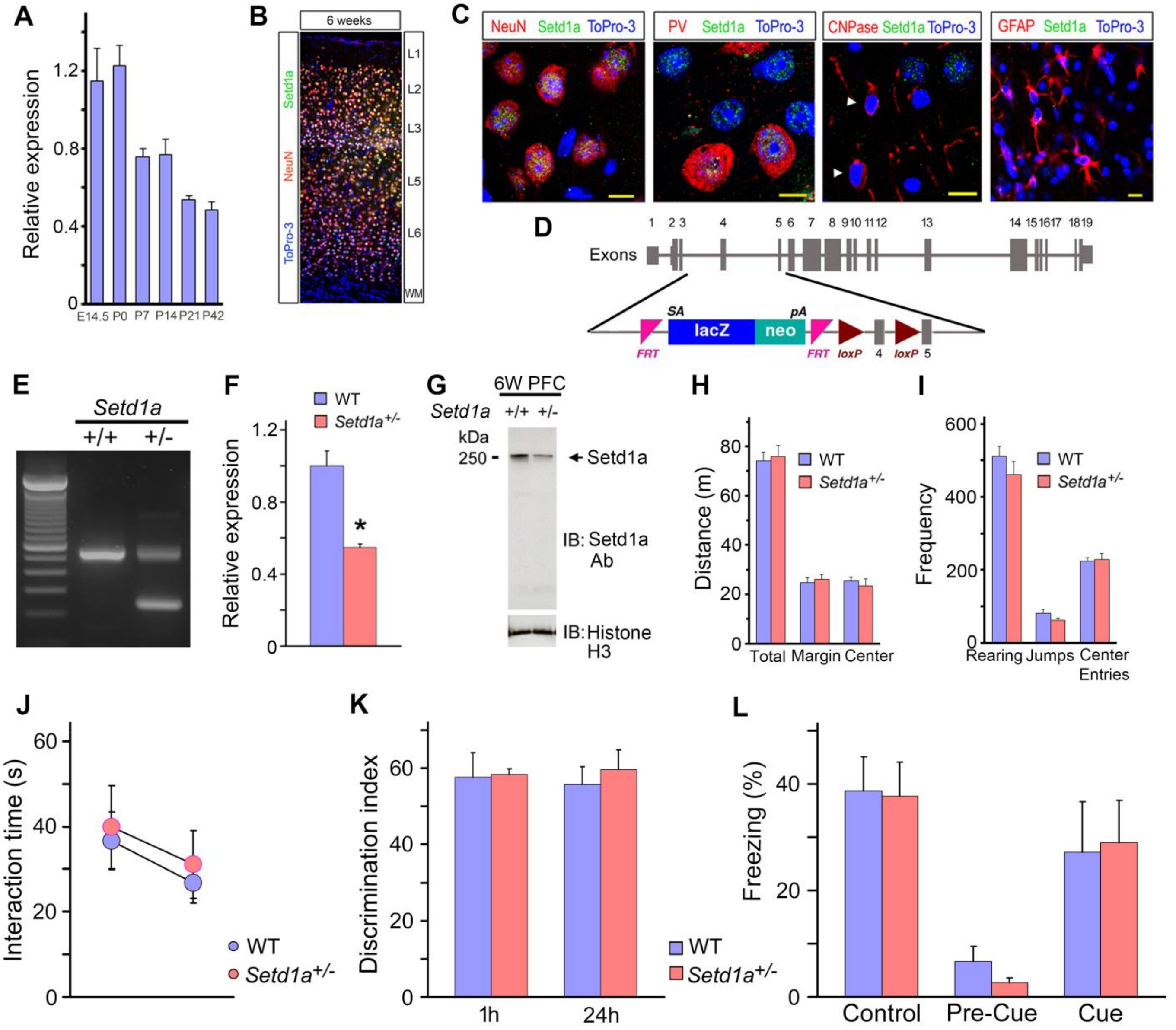

**Supplementary Figure 1: Setd1a expression pattern, molecular and behavioral characterization of *Setd1a*$^{+/-}$ mice** (related to Figure 1)

(A) Developmental expression of *Setd1a* by qRT-PCR in frontal cortex of WT mice (N = 5 each age). Histograms show means and error bars represent s.e.m.

(B) Representative image of Setd1a and NeuN-labeled cells in the prelimbic area from coronal sections of 6 weeks-old brains. WM: white matter.

(C) Higher magnification images from coronal sections of 6 weeks-old brains. Setd1a (*green*) exhibits mixtures of punctate and diffuse fibrillar staining in NeuN$^{+}$ cells (*red* in *left panel*), and PV$^{+}$ interneurons (*red* in *middle left panel*) in medial PFC superficial layers, but neither in CNPase$^{+}$ oligodendrocytes [*red* (*white arrowheads*) in *middle right panel*) in medial PFC superficial layers nor in GFAP$^{+}$ cells (*red* in *right panel*) in corpus callosum. Setd1a localizes to euchromatin (*left* and *middle left panels*), as neither signal overlaps with ToPro-3 (*blue*) bright heterochromatin regions. Scale bar: 10 $\mu$m.

(D) Schematic of the *Setd1a* gene and the targeted locus. The *Setd1a* conditional allele in the *Setd1a*$^{tm1a(EUCOMM)Wtsi}$ mouse line (referred to as *Setd1a*$^{+/-}$) is based on the 'knockout-first' design, a strategy that combines both a reporter-tagged and a conditional mutation. The 'KO-first' allele contains a *LacZ* trapping cassette and a floxed promoter-driven neo cassette inserted into intron 3 of the gene, disrupting gene function.

(E) PCR genotyping showing the expected bands for *Setd1a*$^{+/-}$.

(F) Expression of *Setd1a* by qRT-PCR in medial pre-frontal cortex (mPFC) of *Setd1a*$^{+/-}$ (N = 5) and WT (N = 5) littermate mice at 6-week-old. N, number of individual animals. Data are shown as means ± s.e.m. *$P < 0.05$; **$P < 0.01$. Student's two-tailed *t*-test.

(G) Western blot analysis of Setd1a in the frontal cortex of *Setd1a*$^{+/-}$ and WT littermate mice at 6-week-old.

(H and I) *Setd1a*$^{+/-}$ mice show normal locomotor activity and anxiety: No difference in locomotor activity as assessed by total distance covered between *Setd1a*$^{+/-}$ (N = 11) and WT (N = 10) mice during 60 min exposure to Open Field (H). No difference between genotypes was detected in margin distance vs center distance covered, indicators of anxiety (H). Also, no difference was detected in the number of rearings, jumps or center entries between *Setd1a*$^{+/-}$ (N = 11) and WT (N =10) mice (I). N, number of individual animals (H and I). Data are shown as means ± s.e.m. $P > 0.05$. Student's two-tailed *t*-test.

(J) Intact social memory in *Setd1a* deficient mice. *Setd1a*$^{+/-}$ and WT mice engaged in social interaction with the stimulus juvenile for a similar length of time (N = 8 each genotype, $P > 0.05$). (K) Intact short and long-term Novel Object Recognition memory in *Setd1a* $^{+/-}$ mice. During an 1 h retention test *Setd1a*$^{+/-}$ and WT mice displayed enhanced preference for the novel object to a similar degree (1 h: N = 8 each genotype, $P > 0.05$). Intact long-term Novel Object Recognition memory in *Setd1a*$^{+/-}$ mice. During a 24 h retention test for assessment of long-term recognition memory, *Setd1a*$^{+/-}$ and WT mice displayed comparable preference for the novel object (24 h: N = 8 each genotype, $P > 0.05$).

(L) *Setd1a* deficiency does not affect fear-related associative memory. *Setd1a*$^{+/-}$ and WT mice spent similar percentage of time freezing upon re-exposure to the context in which fear conditioning took place 24 h later. Presentation of the conditioned stimulus along 2 h after re-exposure to the conditioning context led to similar freezing times in *Setd1a*$^{+/-}$ and WT mice (N = 8 each genotype). N, number of independent animals. Data are shown as means ± s.e.m. *$P < 0.05$, **$P < 0.01$. Student's two-tailed *t*-test (K, L) and Two-way ANOVA post hoc pairwise comparisons with Bonferroni correction (J).

**Supplementary Figure 2**

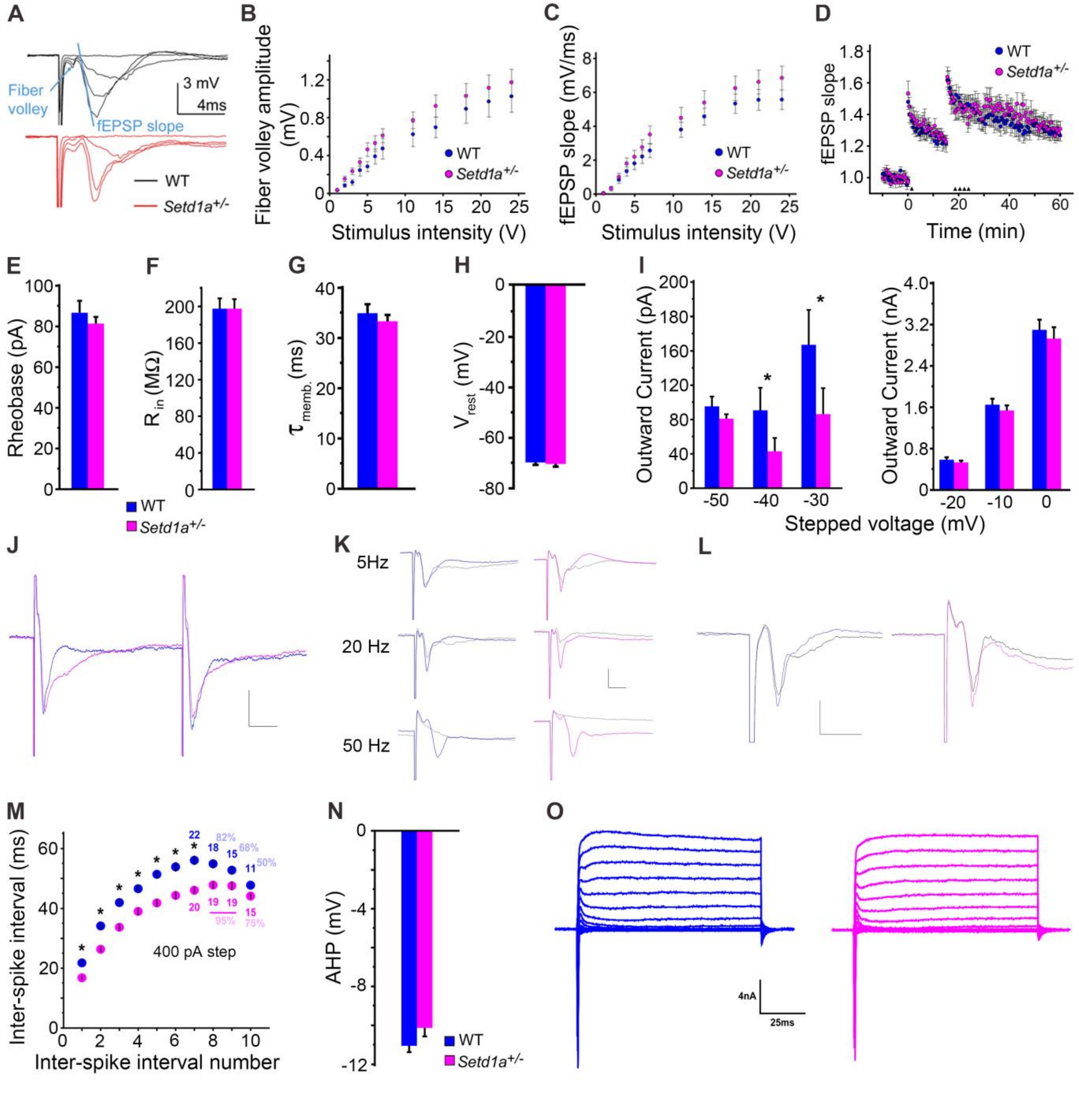

**Supplementary Figure 2: Electrophysiological characterization of *Setd1a*$^{+/-}$ mice** (related to Figure 1)

(A) Sample traces obtained in response to increasing stimulation intensities and showing the fiber volley amplitude (*blue arrow*) as well as the fEPSP initial slope (*blue line*).

(B) Normal afferent volley amplitude in *Setd1a*$^{+/-}$ mice (N = 7, n = 15), compared to WT mice (N = 7, n = 14) (2- way repeated-measures ANOVA, $P > 0.05$).

(C) Normal stimulus-response curve across experiments in *Setd1a*$^{+/-}$ mice (N = 7, n = 25), compared to WT mice (N = 7, n = 26) (2-way repeated-measures ANOVA, $P > 0.05$). WT mice (blue circles); *Setd1a*$^{+/-}$ mice (magenta circles). N, number of individual animals, n, number of brain slices (A-C). Data are shown as means ± s.e.m.

(D) Comparable STP and LTP in *Setd1a*$^{+/-}$ mice overtime. (WT mice: N = 7, n = 12 and *Setd1a*$^{+/-}$ mice: N = 7, n = 11; 2-way repeated-measures ANOVA, $P > 0.05$). STP was induced at the end of the first 50 Hz train stimulation (1 upwards filled arrowhead), and LTP was induced after the stimulation of 4 consecutive 50 Hz trains (4 upwards filled arrowheads). N, number of independent animals. n, number of brain slices. Data are shown as means ± s.e.m. *$P < 0.05$, **$P < 0.01$.

(E) Minimal current necessary to elicit an action potential (rheobase) does not differ between genotypes. Values represent mean ± s.e.m.. (WT, *Setd1a*$^{+/-}$: n = 22, n = 20; unpaired t-test, $P = 0.87$)

(F) Neuronal input resistances ($R_{in}$) were not different between genotypes. Values represent mean ± s.e.m. (WT, *Setd1a*$^{+/-}$: n = 22, n = 20; unpaired t-test, $P = 0.72$)

(G) The neuronal membrane time constant did not differ between genotypes. Values represent mean ± s.e.m. (WT, *Setd1a*$^{+/-}$: n = 22, n = 20; unpaired t-test, $P = 0.41$)

(H) Resting membrane potentials ($V_{rest}$) were not different between genotypes. Values represent mean ± s.e.m. (WT, *Setd1a*$^{+/-}$: n = 23, n = 21; unpaired t-test, $P = 0.66$)

(I) Outward currents evoked by voltage-steps near AP threshold voltages (from -70mV) were reduced in Setd1a+/- mice. Values represent mean ± s.e.m. (WT, Setd1a+/-: n = 30, n = 33; unpaired t-test, -40mV, P = 0.044; -30mV, P = 0.038).

(J) Representative traces showing PPR responses at 50ms, WT neurons (blue trace) and Setd1a$^{+/-}$ neurons (magenta trace). Scale bar: 1 mV, 10 ms. (Related to Fig. 1G)

(K) Representative traces showing STD responses at 5, 20 and 50Hz WT neurons (1st sweep showing in blue and 40th sweep showing in grey) and *Setd1a*$^{+/-}$ neurons (1st sweep showing in magenta and 40th sweep showing in grey). Scale bar: 1 mV, 5 ms. (Related to Fig. 1H)

(L) Representative traces showing LTP responses, WT neurons (baseline trace showing in blue and 1st sweep after 4 consecutive stimulations showing in grey) and Setd1a+/- neurons (baseline trace showing in magenta and 1st sweep after 4 consecutive stimulations showing in grey). Scale bar: 1 mV, 5ms. (Related to Fig. S1D)

(M) Summary data of inter-spike intervals extracted from same neuron recording set shown in Fig. 1I. Values represent mean ± s.e.m. WT, blue; Setd1a$^{+/-}$, magenta. (WT, Setd1a$^{+/-}$: n=22, n=20; unpaired t-test, * $p < 0.05$. Attrition of n-number and percent of recordings included at later inter-spike intervals indicated within panel). (Related to Fig. 1I)

(N) Summary data of AP afterhyperpolarization (AHP) extracted from same neuron recording set shown in Fig. 1I. Values represent mean ± s.e.m. WT, blue; Setd1a$^{+/-}$, magenta. (WT, Setd1a$^{+/-}$: n=22, n=20; unpaired t-test, p=0.063). (Related to Fig. 1I)

(O) Representative current responses to voltage-steps showing activation of fast, transient inward currents and non-inactivating potassium currents. Cells held at -70, stepped from -100mV to +50mV. WT, blue; Setd1a$^{+/-}$, magenta. Scale bar 4nA, 25ms. (Related to Fig. S1I)

### Supplementary Figure 3

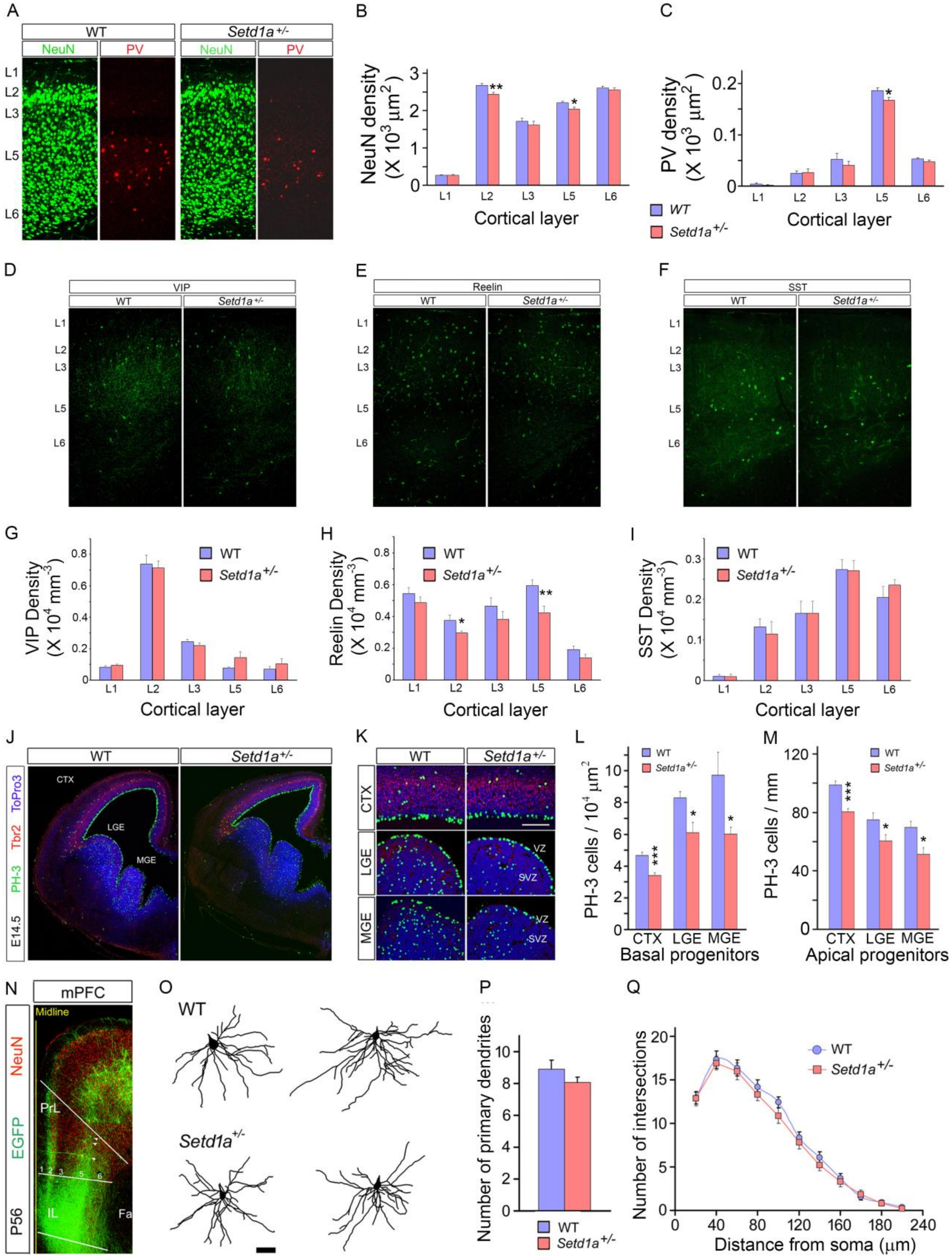

**Supplementary Figure 3. The effect of *Setd1a*-deficiency on cytoarchitectural alterations, proliferation of NSCs at E14.5, and dendritic morphology of pyramidal neurons** (related to Figure 2)

(A-I) Cytoarchitectural changes in the cortex of *Setd1a*$^{+/-}$ mice.

(A) Images of coronal sections of the prelimbic area in medial PFC of 8-week-old WT and *Setd1a*$^{+/-}$ mice showing neuronal cells labeled with NeuN (*green*) and interneurons labeled with PV antibody (*red*). Representative images (A) are composites of more than one image acquired from one brain section under identical scanning parameters.

(B) Histograms indicating a modestly diminished NeuN cell density in L2 (9.0% reduction, $p < 0.01$, N = 6, n = 12 per genotype) as well as in L5 (7.5% reduction, $P < 0.02$, N = 6, n = 12 per genotype) of the PFC (A and B).

(C) PV$^{+}$ cell density in *Setd1a*$^{+/-}$ mice was modestly reduced in L5 (11.5% reduction, $P < 0.05$; N = 5, n =15 per genotype] (A and C).

(D-I) Images of coronal sections of the prelimbic area in medial PFC of 8-week-old WT and *Setd1a*$^{+/-}$ mice depicting different interneuron subpopulations as identified by their immunohistochemical markers (*green*), VIP (D), Reelin (E), and SST (F).

(D-F) Representative images are composites of more than one image acquired from one brain section under identical scanning parameters.

(G-I) Histograms depicting quantitative analysis of the cell density of different interneuron subpopulations, VIP (G), Reelin (H), and SST (I) in the prelimbic area in medial PFC. Reelin$^{+}$ cell density in *Setd1a*$^{+/-}$ mice was modestly reduced in L2 (21.2% reduction, $P < 0.05$; N = 8, n =24 per genotype) and L5 (28.8% reduction, $P < 0.01$; N = 8, n =24 per genotype) (E and H).

(J-M) The effect of *Setd1a*-deficiency on proliferation of NSCs at E14.5

(J) Decreased PH-3 labeling in the telencephalon in *Setd1a*$^{+/-}$ mice during corticogenesis.

(K) Images of coronal sections through the cortex (*top panel*), lateral ganglionic eminence (LGE) (*middle panel*), and medial ganglionic eminence (MGE) (*bottom panel*) at E14.5 showing mitotically active cells labeled with PH-3 in the ventricular zone (VZ) and subventricular zone (SVZ) of WT and *Setd1a*$^{+/-}$ mice.

(L and M) Histograms indicating reduced frequency both of mitotic basal (L, WT: 4.67 ± 0.18 cells/$10^4$ $\mu m^2$, N = 10, n = 30; *Setd1a*$^{+/-}$: 3.39 ± 0.18 cells/$10^4$ $\mu m^2$, N = 9, n = 27, *P* < 0.0001) and apical (M, WT: 98.77 ± 2.56 cells/mm, N = 10, n = 30 ; *Setd1a*$^{+/-}$ : 80.50 ± 2.08 cells/mm , N = 9, n = 27, *P* < 0.0001) NPCs (which give rise to pyramidal neurons) in the cortices of *Setd1a*$^{+/-}$ mice. Similar analysis in the interneuron proliferative zones of the medial ganglionic eminence (MGE) and lateral ganglionic eminence (LGE), showed that basal PH-3$^+$ cells were significantly reduced both in LGE (L, WT: 8.30 ± 0.40 cells/$10^4$ $\mu m^2$, N = 8, n = 16; *Setd1a*$^{+/-}$ : 6.11 ± 0.65 cells/$10^4$ $\mu m^2$ , N = 8, n = 16, *P* < 0.02) and MGE (L, WT: 9.74 ± 1.42 cells/$10^4$ $\mu m^2$, N = 8, n = 16; *Setd1a*$^{+/-}$ : 6.01 ± 0.44 cells/$10^4$ $\mu m^2$ , N = 8, n = 16, *P* < 0.04) of *Setd1a*$^{+/-}$ mice. Similarly, apical PH-3$^+$ progenitors were reduced both in LGE (M, WT: 75.01 ± 4.01 cells/mm, N = 8, n = 16; *Setd1a* $^{+/-}$ : 60.55 ± 4.21 cells/mm , N = 8, n = 16, *P* < 0.04), and MGE (M, WT: 69.60 ± 4.21 cells/mm, N = 8, n = 16; *Setd1a*$^{+/-}$: 51.36 ± 4.27 cells/mm , N = 8, n = 16, *P* < 0.01) of *Setd1a*$^{+/-}$ mice.

N, number of independent animals. n, number of brain slices. Data are shown as means ± s.e.m. **P* < 0.05, ***P* < 0.01.

(N) Representative image of EGFP-labeled neurons in layer 5 (*arrow heads*) in PrL of *Thy1-GFP*$^{+/-}$*:Setd1a*$^{+/-}$ mouse. Numbers indicate cortical layers. PrL: prelimbic area, IL: infralimbic area, Fa: corpus callosum anterior forceps. Images are composites of more than one image acquired from one brain section under identical scanning parameters.

(O-Q) Lack of alterations in dendritic complexity in the medial PFC of mutant mice.

(O) Representative tracings of the basal dendritic trees of L5 pyramidal neurons at the prelimbic area of medial PFC from *Setd1a*$^{+/-}$; *Thy1-GFP*$^{+/–}$ and WT mice.

(P) No alterations in number of primary dendrites in the basal dendritic tree of L5 pyramidal neurons between genotypes (n = 18 cells for *Setd1a*$^{+/-}$; n = 18 cells for WT, *P* = 0.21).

(Q) Sholl analysis of basal dendrite complexity using 20 $\mu m$ concentric circles around the soma (n = 18 cells for *Setd1a*$^{+/-}$; n = 18 cells for WT, *P* > 0.05).

### Supplementary Figure 4

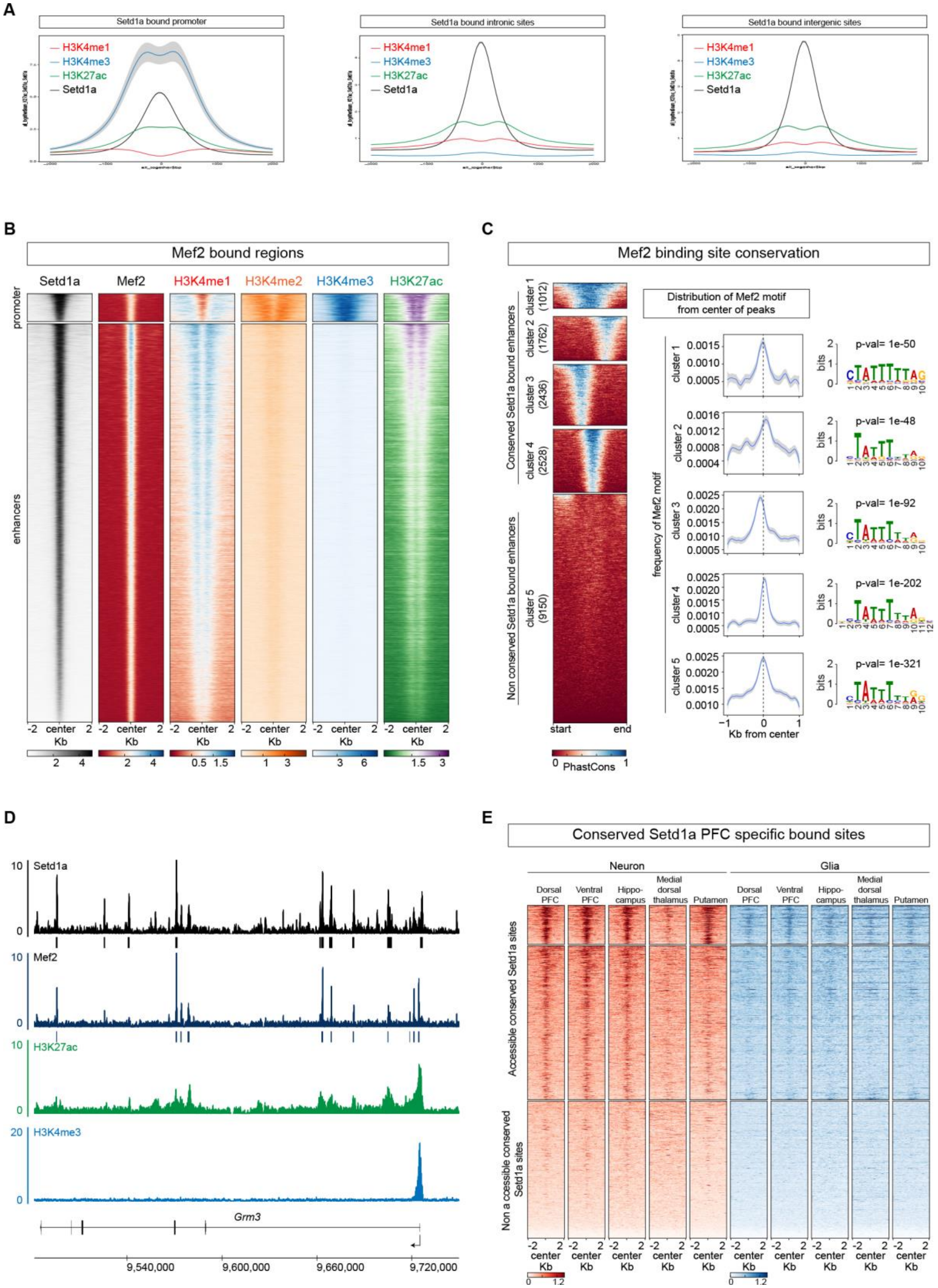

**Supplementary Figure 4: Setd1a and Mef2 co-bind PFC specific regulatory elements** (related to Figure 3)

(A) Distribution of the mean normalized coverage 2 kb around the center of Setd1a peaks, annotated in promoter (*left*), intron (*middle*) and intergenic regions (*right*). Coverage of PFC ChIP-Seq signal for H3K4me1 (*red*), H3K4me3 (*blue*), H3K27ac (*green*) and Setd1a (*black*) are shown. Shading indicates 95% confidence intervals.

(B) Heatmap of the mean normalized coverage for Setd1a (*black*), Mef2 (*dark blue*) and chromatin marks (H3K4me1, *red*; H3K4me2, *orange*; H3K4me3, *light blue*; H3K27ac, *green*) ChIP-Seq of Mef2 peaks. We distinguished Mef2 bound peaks at promoters (*top panel*) (n=2449) marked by H3K4me3, and at enhancers (*bottom panel*) (n=22701) marked by H3K4me1 and H3K27ac.

(C) Heatmap of PhastCons values per base pairs of Setd1a peaks normalized by size (*left*), distribution and motif enrichment of the motif within Setd1a peaks of each cluster (*right*). Five clusters were defined according to their conservation level: one very conserved cluster (*cluster1*), three partially conserved clusters (*cluster2, cluster3, cluster4*) and one poorly conserved cluster (*cluster5*). Each cluster is enriched for Mef2 motifs (*right panels*) and significant p-value is indicated.

(D) The *Grm3* locus is shown as an example of regulatory regions bound by Setd1a and Mef2 in the PFC. In this locus, we saw extensive binding overlap between Setd1a and Mef2 at both the promoter and intragenic enhancers. The normalized reads per genomic coverage for Setd1a (*black*), Mef2 (*dark blue*), H3K27ac (*green*), H3K4me3 (*light blue*) are shown. Black and dark blue boxes, below Setd1a and Mef2 tracks respectively, represent significant peaks that passed all quality checks (see methods).

(E) Heatmap of ATAC-Seq coverage 2 kb around the center of the most conserved PFC specific Setd1a bound enhancers (see Figure 3C), between mouse and human (megablast, see Methods and Data Table 1), from different regions (dorsal PFC, ventral PFC, Hippocampus, Medial dorsal Thalamus and Putamen) of postmortem human brain from sorted neuronal (*red*) and glia (*blue*) cells.

**Supplementary Figure 5**

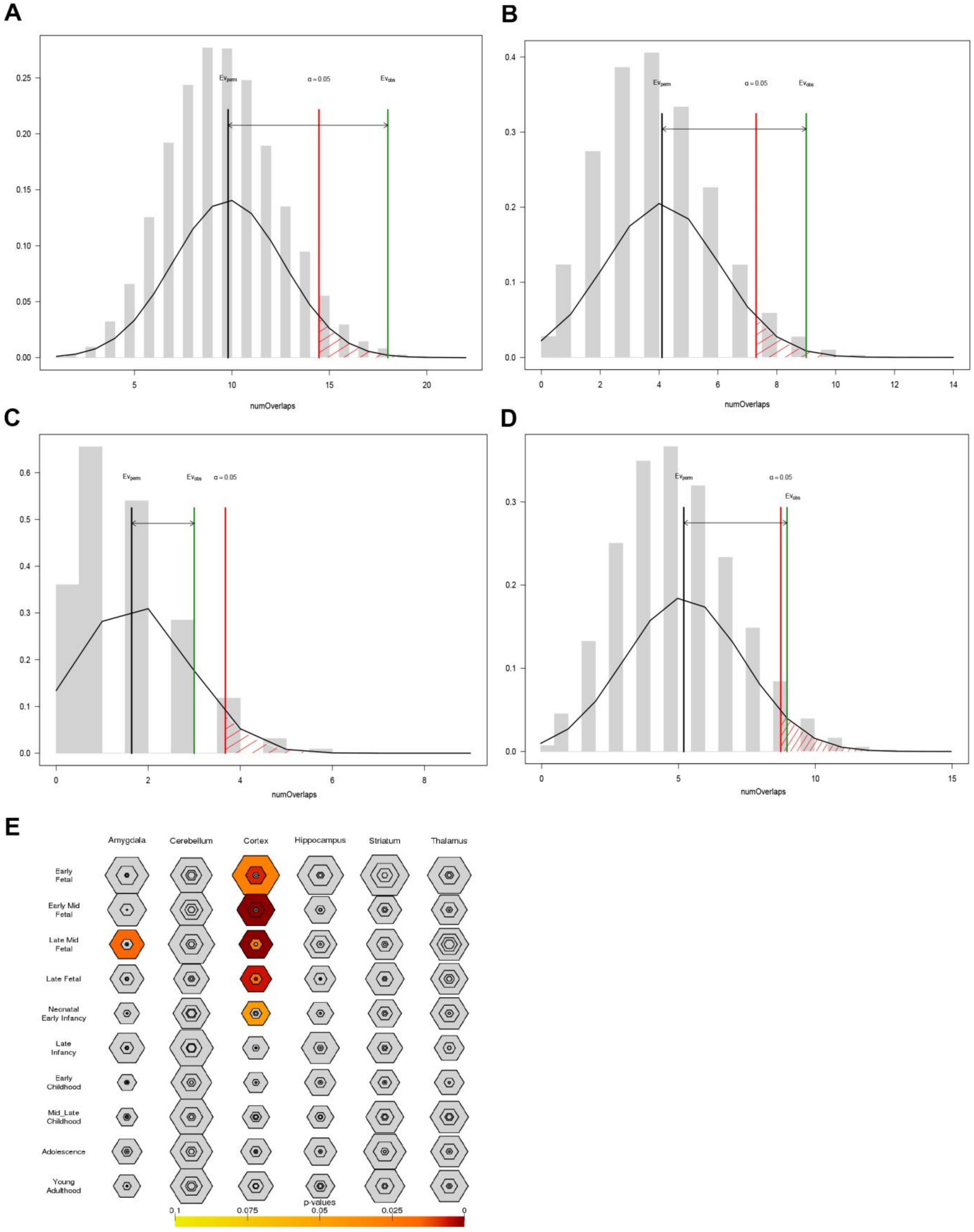

**Supplementary Figure 5: Overlap between genome-wide significant SCZ signals and conserved (by the megablast method) chromatin accessible Setd1a ChIP-Seq peaks** (related to Figure 4)

(A) All Setd1a peaks vs SCZ GWAS intervals. *P* = 0.0055, Z-score = 2.9.

(B) Setd1a Intronic peaks vs SCZ GWAS intervals. *P* = 0.021, Z-score = 2.5.

(C) Setd1a exonic and UTR peaks vs SCZ-GWAS intervals. *P* = 0.222, Z-score = 1.1.

(D) Setd1a Intergenic peaks vs SCZ-GWAS intervals. *P* = 0.074, Z-score = 1.75.

All tests use permutations (n=20,000, randomization method = circularRandomizeRegions) to determine if there are significant overlaps between conserved Setd1a ChIP-Seq peaks and previously identified SCZ-GWAS loci (Schizophrenia Working Group of the Psychiatric Genomics, 2014). X axis represents the number of overlaps, Y axis represents the density of expected number of overlaps determined by permutation. EVperm: Expected average number of overlaps by permutation (*black line*), EVobs: actual observed number of overlaps (*green line*). α, permutation *P* value threshold = 0.05 (*red line*). SCZ GWAS: the independent schizophrenia GWAS intervals calculated in the original publication (Schizophrenia Working Group of the Psychiatric Genomics, 2014).

(E) Spatiotemporal expression pattern of target genes within conserved and accessible Setd1a peaks that overlap with the mutation intolerant genes hit by LoF DNMs in NDD show prenatally-biased expression in various brain regions.

### Supplementary Figure 6

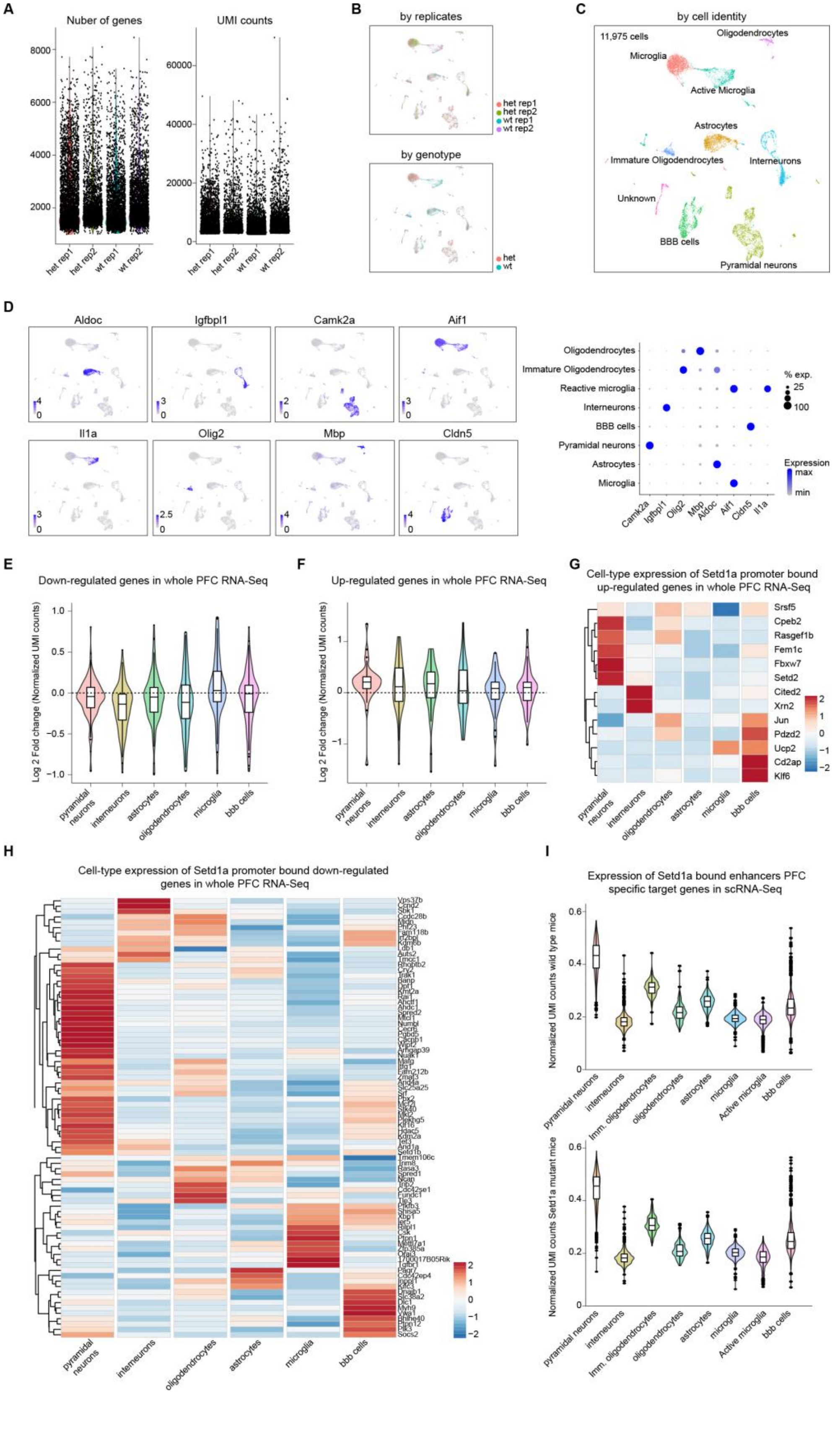

**Supplementary Figure 6: scRNA-Seq quality checks and analysis of cell-type expression of Setd1a target genes** (related to Figure 5)

(A) Number of genes (*left panel*) and UMI counts (*right panel*) per replicate of scRNA-Seq performed on murine PFC (2 biological replicates per wild type and Setd1a mutant mice).

(B) UMAP representation of 11,975 single cell transcriptomes from the PFC of wild-type (5,944) and *Setd1a*$^{+/-}$ (6,031) mice colored by replicates (*top panel,* PFC from Setd1a heterozygous mice (het) replicate 1 (red) and 2 (green), PFC from wild type mice replicate 1 (blue) and 2 (purple)) and by genotype (Setd1a heterozygous mice (het, red), PFC from wild type mice (blue)) (*bottom panel*).

(C) UMAP representation of 11,975 single cell transcriptomes from the PFC of wild-type and *Setd1a*$^{+/-}$ mice highlighting the different clustered cell-types.

(D) Expression of several known cell type marker genes for neuronal and non-neurons cells (*left panels*): *CamK2a* (pyramidal neurons), *Igfbpl1* (interneurons), *Aldoc* (astrocytes), *Aif1* (microglia), *Olig2* (oligodendrocytes), *Mbp* (mature oligodendrocytes), *Cldn5* (endothelial cells), *Il1a* (reactive microglia). Dot-plot showing the expression of neuronal and glial markers (*right panels*).

(E-F) Expression differences within the scRNA-Seq dataset of the down-regulated (E) and up-regulated (F) genes identified by whole PFC RNA-Seq (see Figure 3A). The log2 fold change ratio of Setd1a$^{+/-}$/WT normalized UMI count per cell-type is shown. Biological replicates per genotype were pooled together.

(G-H) Heatmaps of the expression of up-regulated (G) and down-regulated (H) genes with Setd1a bound promoters (as defined in Figure 5D), across the different cell types identified by scRNA-Seq. Heatmaps show the normalized UMI counts per gene. All scRNA-Seq datasets were pooled together.

(I) Average expression of all genes with enhancers, but not promoters, bound by Setd1a specifically in the PFC (see Figure 3C), per genotype (WT, *top panel*; *Setd1a*$^{+/-}$, *bottom panel*). We observed that the average expression of these genes in pyramidal neurons is significantly higher than any other cell-types (Wilcox test < 0.001).

**Supplementary Figure 7**

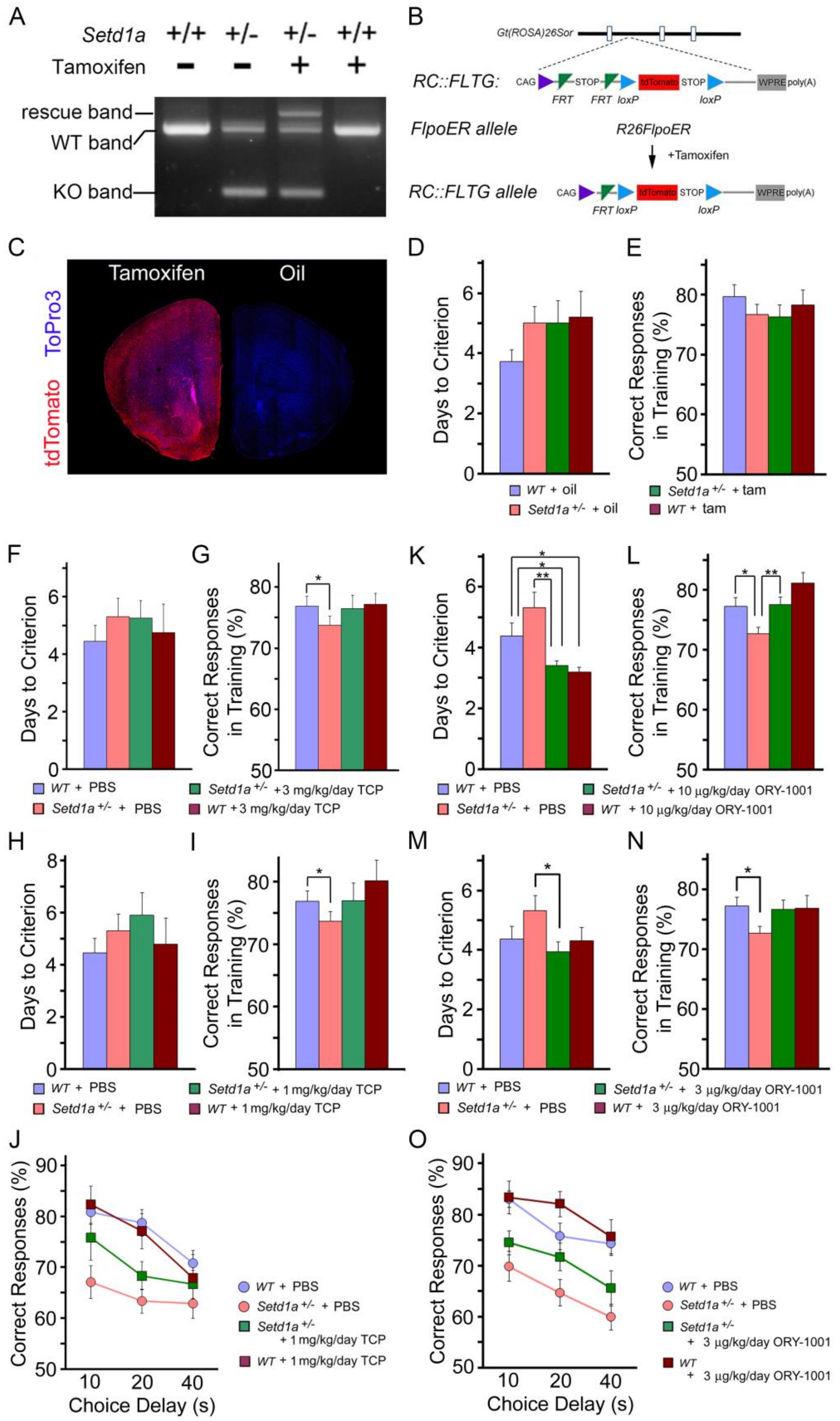

**Supplementary Figure 7: WM performance following *Setd1a* re-activation and pharmacological inhibition of LSD1 in adulthood** (related to Figure 6 and Figure 7)

(A-D) Activation of *Setd1a* expression in adulthood and effects on WM performance.

(A) PCR genotyping showing the expected bands (KO, rescue and WT) generated by tamoxifen inducible Flp-mediated recombination at the targeted *Setd1a* genomic locus

(B) Schematic representation of tamoxifen-inducible reporter expression strategy. The Flp responsive fluorescent indicator allele, *RC:FLTG* was previously generated by targeting intron 1 of the *Gt(ROSA)26Sor* locus. FRT flanked transcriptional stop cassettes in *RC::FLTG* prevent expression of tdTomato in the absence of Flp activity. Tamoxifen-inducible Flpo expression excises stop cassettes and leads to tdTomato expression in $R26^{FlpoER+/-}$ *:RC: :*$FLTG^{+/-}$ mice.

(C) Tamoxifen-inducible Flp strategy leads to broad reporter expression. Coronal sections from $R26^{FlpoER+/-}$*:RC::*$FLTG^{+/-}$ mice after feeding with tamoxifen (*left*) or corn oil (*right*). Results show widespread induction of tdTomato reporter expression after tamoxifen–induced Flp activation, but not in the absence of Flp activity (corn oil).

(D) Days taken to criterion performance (three consecutive days of > 70% choice accuracy) during training on the delayed non-match to sample T-maze task following activation of Setd1a expression in adulthood.

(E) Correct responses during training on the delayed non-match to sample T-maze task following activation of Setd1a expression in adulthood.

(F and G) TCP administration in adulthood and effects on WM performance. Days taken to criterion performance (three consecutive days of > 70% choice accuracy) (F) and mean percentage of correct responses for WT and $Setd1a^{+/-}$ mice (G) during training on the delayed non-match to sample T-maze task following 3 mg/kg/day TCP or vehicle treatment. *$P < 0.05$.

(H-J) Low concentration (1 mg/kg/day) of TCP does not rescue WM performance in the delayed non-match to place task. Days taken to criterion performance (three consecutive days of > 70% choice accuracy) (H), and mean percentage of correct responses during training (I) were not affected. The impairment in the performance of vehicle-treated mutant mice over vehicle treated WT control littermates during testing with increasing time delays was not completely reversed by 1 mg/kg/day TCP treatment (J). *$P$, 0.05. **$P$, 0.01, ***$P$, 0.001, Data are mean ± s.e.m (*Setd1a*$^{+/-}$ + PBS: N = 20, *Setd1a*$^{+/-}$ + TCP: N = 18, WT + PBS: N = 20, WT + TCP: N = 20).

(K and L) WM performance following administration of 10 μg/kg/day ORY-1001 in adulthood. Days to criterion (K) and percentage of correct choices (L) during training phase were improved in both ORY-1001-treated *Setd1a*$^{+/-}$ and WT mice. *$P$, 0.05. **$P$, 0.01, ***$P$, 0.001, Data are mean ± s.e.m (*Setd1a*$^{+/-}$ + PBS: N = 21, *Setd1a*$^{+/-}$ + ORY-1001: N = 15, WT + PBS: N = 23, WT + ORY-1001: N = 15).

(M-O) Low concentration (3 μg/kg/day) of ORY-1001 does not rescue WM performance in the delayed non-match to place task. Shown are days taken to criterion performance (three consecutive days of > 70% choice accuracy) during training (M) and mean percentage of correct responses during training (N). An impairment in the performance of vehicle-treated mutant mice over vehicle treated WT control littermates during testing with increasing time delays was not effectively reversed by 3 μg/kg/day ORY-1001 treatment (O) *$P$, 0.05. **$P$, 0.01, ***$P$, 0.001, Data are mean ± s.e.m (WT + PBS: N = 23, *Setd1a*$^{+/-}$ + PBS: N = 21, *Setd1a*$^{+/-}$ + ORY-1001: N = 15, WT + ORY-1001: N = 15).

**Table S1**

| GWAS Category | # of signal peaks | Permutation P value (n = 20,000) | # of Overlaps | Z scores |
|---|---|---|---|---|
| SCZ_PGC | 105 | 0.006 | 18 | 2.90 |
| Blood_metabolite_levels | 160 | 0.006 | 1 | -2.03 |
| Body_mass_index | 131 | 0.546 | 7 | -0.13 |
| Bone_mineral_density | 78 | 0.506 | 4 | 0.11 |
| Breast_cancer | 80 | 0.442 | 2 | -0.53 |
| Cholesterol_total | 82 | 0.009 | 0 | -2.22 |
| Crohns_disease | 127 | 0.135 | 3 | -1.28 |
| HDL_cholesterol | 98 | 0.525 | 4 | -0.28 |
| inflammatory_bowel_disease | 113 | 0.038 | 1 | -1.82 |
| LDL_cholesterol | 75 | 0.158 | 1 | -1.19 |
| Menarche | 129 | 0.429 | 7 | -0.42 |
| Prostate_cancer | 100 | 0.53 | 5 | 0.05 |
| QT_interval | 78 | 0.503 | 4 | 0.07 |
| Rheumatoid_arthritis | 101 | 0.381 | 4 | -0.60 |
| Type_2_diabetes | 86 | 0.546 | 4 | 0.03 |

**Supplementary Table 1: Overlap between conserved accessible Setd1a Chip-Seq peaks and GWAS signals of various complex disorders** (related to Figure 4)

**Table S2**

| GWAS Category | # of signal peaks | Permutation P value (n = 20,000) | # of Overlaps | Z scores |
|---|---|---|---|---|
| Alzheimer | 51 | 0.379 | 1 | -0.754 |
| ALS | 18 | 0.564 | 0 | -0.714 |
| Autism | 20 | 0.292 | 1 | -0.978 |
| Parkinson | 43 | 0.108 | 5 | 1.555 |

**Supplementary Table 2: Overlap between conserved accessible Setd1a Chip-Seq peaks and GWAS signals of various CNS diseases** (related to Figure 4)

**Table S3**

| GWAS Category | Adj P value | # of genes in GWAS intervals |
|---|---|---|
| SCZ_PGC | 0.008 | 15 |
| Body mass index | 0.569 | 7 |
| Cholesterol total | 0.976 | 2 |
| Blood_metabolite_levels | 1.000 | 2 |
| Blood_metabolite_ratios | 1.000 | 0 |
| Bone_mineral_density | 0.652 | 4 |
| Breast_cancer | 0.910 | 3 |
| HDL_cholesterol | 0.513 | 5 |
| height | 0.988 | 11 |
| inflammatory_bowel_disease | 0.952 | 3 |
| Inflammatory_skin_disease | 0.422 | 4 |
| LDL_cholesterol | 1.000 | 1 |
| Menarche | 0.451 | 8 |
| Metabolite_levels | 0.790 | 3 |
| Multiple_sclerosis | 0.918 | 2 |
| Platelet_count | 0.249 | 5 |
| Prostate_cancer | 0.578 | 4 |
| QT_interval | 0.571 | 4 |
| Red_blood_cell_traits | 0.765 | 2 |
| Rheumatoid_arthritis | 0.907 | 3 |
| Type_2_diabetes | 0.903 | 3 |

**Supplementary Table 3: Overlap between conserved accessible Setd1a target genes and GWAS signals of various complex disorders** (related to Figure 4)

**Table S4**

| INTERVAL | GENE_LOC | GENE_ID | GENE_SYMBOL | GENE_DESC |
|---|---|---|---|---|
| int11:chr1:98375391..98559093 | chr1:97543299..98386615 | 1806 | DPYD | Dihydropyrimidine_dehydrogenase |
| int62:chr2:198148191..198835702 | chr2:197851385..198175521 | 91526 | ANKRD44 | Ankyrin_repeat_domain_44 |
| int63:chr2:200162425..200314206 | chr2:200134222..200335989 | 23314 | SATB2 | SATB_homeobox_2 |
| int77:chr3:2532788..2561691 | chr3:2142246..3099645 | 152330 | CNTN4 | Contactin_4 |
| int84:chr4:170357792..170646003 | chr4:170314420..170533778 | 4750 | NEK1 | NIMA_(never_in_mitosis_gene_a)-related_kinase_1 |
| int84:chr4:170357792..170646003 | chr4:170541721..170642157 | 1182 | CLCN3 | Chloride_channel_3 |
| int85:chr4:176851045..176875795 | chr4:176554087..176923648 | 2823 | GPM6A | Glycoprotein_M6A |
| int93:chr5:45291514..45393754 | chr5:45259351..45696220 | 348980 | HCN1 | Hyperpolarization_activated_cyclic_nucleotide-gated_potassium_channel_1 |
| int114:chr7:86403263..86459347 | chr7:86273229..86494192 | 2913 | GRM3 | Glutamate_receptor,_metabotropic_3 |
| int107:chr7:104594253..105063372 | chr7:104756822..105029341 | 6733 | SRPK2 | SRSF_protein_kinase_2 |
| int109:chr7:110843795..111180544 | chr7:110303109..111202347 | 83943 | IMMP2L | IMP2_inner_mitochondrial_membrane_peptidase-like_(S._cerevisiae) |
| int111:chr7:137042224..137085250 | chr7:137074384..137531609 | 9162 | DGKI | Diacylglycerol_kinase,_iota |
| int119:chr8:4177231..4192528 | chr8:2792874..4852328 | 64478 | CSMD1 | CUB_and_Sushi_multiple_domains_1 |
| int17,int18:chr10:18680963..18770063 | chr10:18429605..18830688 | 783 | CACNB2 | Calcium_channel,_voltage-dependent,_beta_2_subunit |
| int54,int55:chr18:52937841..53200117 | chr18:52889561..53255860 | 6925 | TCF4 | Transcription_factor_4 |

**Supplementary Table 4: High-confidence Setd1a target genes within the SCZ-GWAS loci** (related to Figure 4)

**Table S5**

| GWAS Category | Adj P value | # of genes in GWAS intervals |
|---|---|---|
| Alzheimer | 1.000 | 1 |
| ALS | 0.274 | 2 |
| Autism | 0.015 | 2 |
| Parkinson | 1.000 | 1 |

**Supplementary Table 5: Overlap between conserved accessible Setd1a target genes and GWAS signals of various CNS diseases** (related to Figure 4)

**Table S6**

| | # of genes tested | *Adj P* value | # of genes in SCZ GWAS intervals |
|---|---|---|---|
| Setd1a bound targets | 430 | 0.008 | 15 |
| Setd1a and Mef2 co-bound targets | 367 | 0.003 | 15 |
| Mef2 only bound targets | 60 | 0.033 | 4 |

**Supplementary Table 6: Enrichment of Setd1a and Mef2 bound targets in the SCZ** (related to Figure 4)
